# A high throughput investigation of the binding specificity of carbohydrate-binding modules for synthetic and natural polymers

**DOI:** 10.1101/2025.02.25.640029

**Authors:** Andrew Philip Rennison, Jaime Fernandez-Macgregor, Julie Melot, Fabien Durbesson, Tobias Tandrup, Peter Westh, Renaud Vincentelli, Marie Sofie Møller

## Abstract

Carbohydrate-binding modules (CBMs) are non-catalytic domains that enhance enzyme binding to substrates, crucial for polysaccharide degradation. Type A CBMs also show potential for engineering plastic-degrading enzymes due to their high affinity for synthetic polymers. This study presents a high-throughput screening pipeline for characterizing the affinity and specificity of Type A CBMs to the synthetic polymers polyethylene terephthalate (PET), polystyrene (PS), and polyethylene (PE), and the natural polysaccharides cellulose, chitin, and starch. Approximately 800 CBMs from the families CBM2, CBM3, CBM10, and CBM64 were expressed as enhanced green fluorescent protein (EGFP)-fusion proteins and tested for binding using a modified holdup assay, which could produce up to 10,000 data points per day. The screening identified approximately 150 binders for PET and PE, around 250 for PS, and demonstrated family-specific binding patterns for avicel, chitin, and starch. Distinct patterns of substrate affinity and specificity were observed, which allowed for rationalizations of binding at the structural and phylogenetic levels. To demonstrate practical utility, CBMs with high PET affinity were fused to the PET hydrolase LCC^ICCG^, enhancing enzymatic activity on crystalline PET powder by around 5-fold. Importantly, these CBM-enzyme fusions mitigated competitive binding to inert plastic impurities, improving performance in mixed plastic assays. This work significantly expands the known repertoire of CBMs capable of binding synthetic polymers, advances our understanding of CBM-substrate interactions, and provides knowledge for engineering enzymes to tackle plastic pollution, particularly in municipal solid waste where mixed plastics pose significant challenges.

**Significance Statement:** Improper recycling of plastic poses a severe environmental threat, with synthetic polymers persisting in ecosystems for decades. Enzymatic recycling of plastic offers a promising solution, with optimization of the enzymes and processes ongoing. To expand the toolkit for engineering enzymes via substrate-binding module fusion, we developed a high-throughput screening platform to identify carbohydrate-binding modules (CBMs) that selectively bind synthetic plastics such as PET, polystyrene, and polyethylene, as well as natural polysaccharides. By screening 797 CBMs, we discovered binding domains with distinct specificities, enabling the design of enzyme fusions that enhance plastic degradation, even in mixed plastic environments. This approach provides insights into protein-substrate interactions that can be leveraged for both waste management and bioengineering innovations.

## Introduction

Plastic pollution is a prominent environmental issue, driven by the rise in single-use plastics and inadequate disposal systems. This has led to widespread plastic dispersal in the biosphere (1), causing numerous adverse effects (2, 3). The projection that the ocean will contain more plastic waste than fish by 2050 (4) underscores the urgent need to address existing pollution and prevent future waste. Additionally, over 98% of plastic packaging is made from virgin fossil feedstocks, with 6% of global oil consumption going to plastic production (4), highlighting the need for sustainable recycling practices.

Recycling plastic waste and creating a closed-loop economy are often proposed to mitigate the environmental impact of plastics. Generally, plastic recycling can be divided into two types, chemical/physical (5) and enzymatic (6) methods. However, both have issues. Chemical/physical methods are energetically expensive or use polluting reagents like methanol (7), often reducing the quality of recycled polymers (8). Enzymatic recycling, though much less developed, is currently limited to polyethylene terephthalate (PET) degradation, and is not yet industrialized (9, 10). Both recycling pathways require sorting mixed plastic waste into constituent polymers, adding costs and environmental impacts. Crucially, neither can reduce existing plastic waste without collection schemes. Therefore, an *in situ* solution is also needed to remediate plastic pollution.

Numerous studies have identified fungal, bacterial, and algae species acting on various plastics (11). Bacterial degradation of PET has been well reported since the discovery of *Ideonella sakaiensis* in 2016, which can use PET as its sole carbon source (12). Significant degradation of polystyrene (PS) and polyethylene (PE) by bacterial isolates has also been shown (13, 14). Many fungal isolates from polluted environments, particularly from the genus *Aspergillus*, have demonstrated activity on these plastics (15, 16). Fungi, known for their role in the turnover of recalcitrant biomass and production of industrially relevant enzymes for bioprocessing recalcitrant material, secrete enzymes like laccases, peroxidases, and cutinases that act on lignin and complex biopolymers as well as chemically similar polymers such as PE, PET, and PS (17–19). However, enzymatic degradation of PET is much more developed than other plastics. The PAZy database (www.pazy.eu), which catalogs biochemically characterized enzymes, currently lists over 100 PET hydrolases, two non-specific oxidase classes of PE degraders, and no enzymes for PS degradation (20).

In nature, many enzymes involved in bioprocessing natural polymers are modular proteins, including carbohydrate-binding modules (CBMs) besides the catalytic domain. These non-catalytic CBMs can enhance enzyme activity by increasing affinity or enzyme concentration at the substrate. They can also increase enzyme specificity by directing it to a particular polymer within a complex polysaccharide environment such as the plant cell wall (21). Several CBMs that bind to plastics have been identified (22–24), and studies have shown their potential in engineering plastic-degrading enzymes, primarily focusing on PET degradation (25–27). These CBMs are classified as Type A, characterized by a planar binding surface that allows them to bind to insoluble targets (21), and are mainly grouped into CBM families 2 and 3 (CBM2 and CBM3) in the carbohydrate-active enzymes (CAZy) database (www.cazy.org) (28).

While industrial scale enzymatic degradation of pure PET streams may not require a CBM (24), municipal solid waste often comprises mixed plastics like PET, PS, and PE (29). Enzyme cocktails for degrading mixed plastics will likely need substrate-binding domains with specificity for each plastic substrate. Engineering fungal or bacterial strains for *in situ* biodegradation of plastic pollution may also require binding domains, due to the lower substrate concentration compared to industrial conditions. Substrate-binding domains that do not bind to environmentally abundant polysaccharides like cellulose or starch would be beneficial in these conditions.

The binding of CBMs to polysaccharides and their impact on enzymatic activity have been studied for decades. However, many studies have been biased towards polysaccharides that the associated catalytic domains act upon, and systematic studies testing CBMs against various polysaccharides are rare. Some studies screen multiple CBMs from different families on one substrate or test various substrates with CBMs from one family (30, 31). These limited studies do not provide sufficient data to predict binding affinity and selectivity based on sequence, structure, or phylogeny. It has been proposed that the specificity of Type A CBMs is influenced by the spacing between aromatic residues in the binding plane or their relative angles (32, 33). However, these proposals are often based on testing of individual modules and may not apply across the entire family. Many Type A CBM families bind both cellulose and chitin, with a distinction often made between these and starch-binding domains (SBDs). Given the similarity between the surfaces of starch granules and crystalline cellulose (34, 35), some promiscuity in binding could be expected. Understanding CBM specificity for these highly abundant polysaccharides would not only aid in developing enzymes for bioremediation of plastic waste but also enhance our knowledge of biomass turnover on Earth.

In this study, we developed a high-throughput (HTP) screening pipeline (Fig. 1A) for Type A CBMs binding to synthetic polymers (PET, PS, and LDPE) and polysaccharides (cellulose, chitin, and starch). Approximately 1,200 proteins from CBM families 2, 3, 10, and 64 were rationally selected, designed as fusions with an enhanced green fluorescent protein (EGFP), and expressed and purified in an HTP manner. These proteins were analyzed using an adapted holdup (HU) assay (36, 37) (Fig. 1B), with binding intensities (BI) calculated for each substrate (Fig. 1C). CBMs that specifically bound to a given plastic were then fused to the PET-degrading enzyme LCC^ICCG^ (9, 10), and the selectivity of these fusion enzymes was determined in mixed substrate assays.

**Fig. 1.**
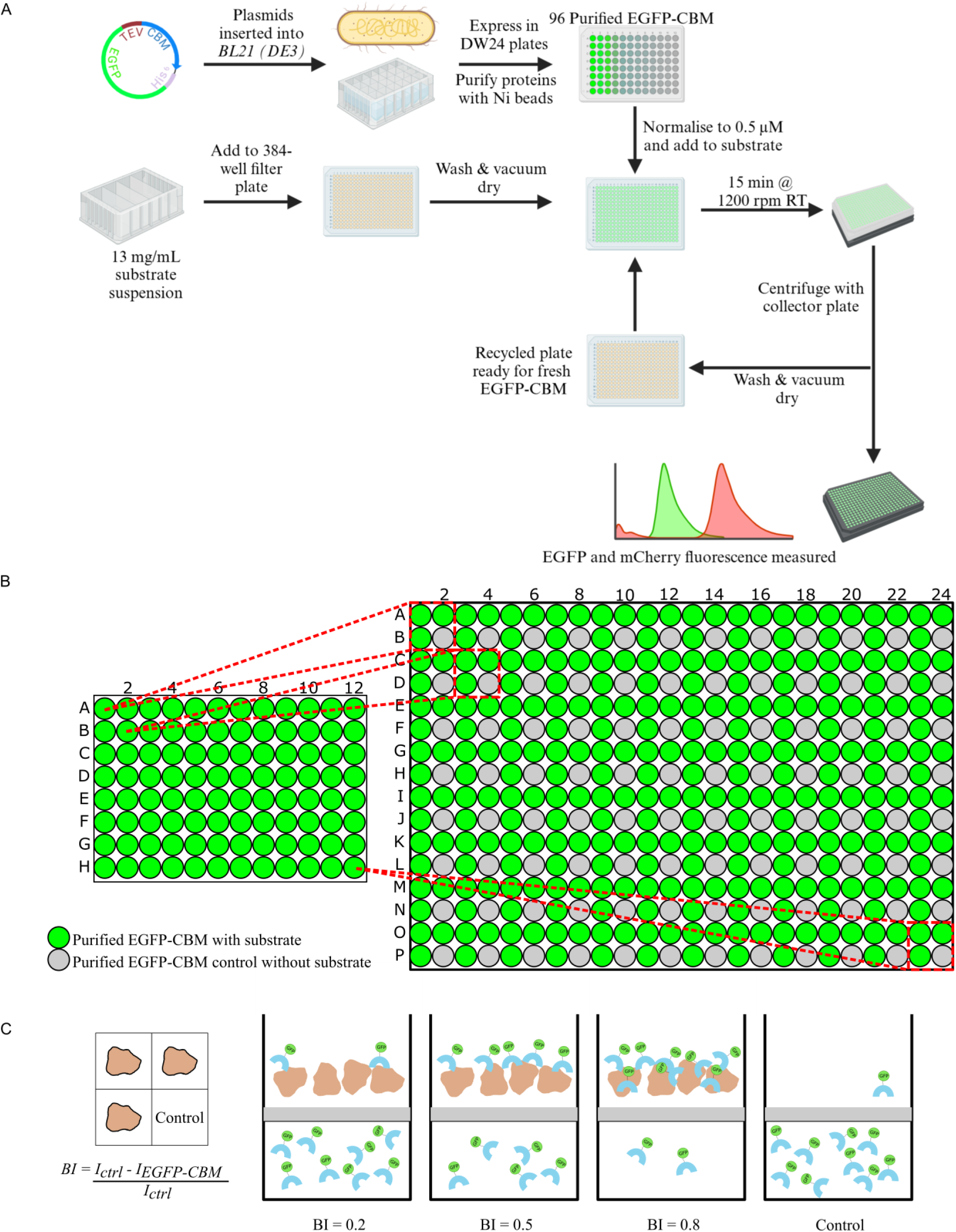
Setup for the high throughput (HTP) holdup assay. (*A*) The process flow of HTP expression, purification, and binding analysis by the holdup assay. (*B*) The setup of the 384-well filter plates used in the assays. The purified EGFP-CBM is added from the 96 well plate, to the 384 well plate in the pattern shown. The bottom right well of each group of four has no substrate added, and is used as the control well. (*C*) The conceptual framework of the holdup assay. Substrate is added to the top of the filter plate, followed by the protein, which binds to the substrate at varying levels. Unbound protein is then removed by centrifugation and collected in a 384 black plate; it is detected by the fluorescent signal from the EGFP module. BI values are calculated as a fraction of the bound total, compared to the control wells.

## Results

### HTP Holdup Assays to Identify Binding Profiles of CBMs

The results and discussion of the selection and expression of the EGFP-CBM proteins can be found in the supporting information. Out of the 1097 EGFP-CBM proteins initially selected for expression, 797 yielded concentrations above 0.5 µM after purification, making them suitable for the HTP holdup assays. Details on the design and validation of the holdup assay can be found in the *SI Appendix* (Supporting information text). BI values were determined for each CBM against the polymers PET, PS, and LDPE, as well as the polysaccharides avicel, chitin, and starch. Constructs with BI values below that of EGFP alone (*SI Appendix,* Table 1) on a given substrate were considered non-binders. Histograms depicting the binding intensities for all binders on each substrate are shown in Fig. 2, with BI values available in *Dataset S1*. The holdup assay identified 727 CBMs that bound to at least one of the six substrates, equating to a 91.2 % hit rate. The raw data and results plotted for each of the 384-well filter plates are provided in *Dataset S2*.

**Fig. 2.**
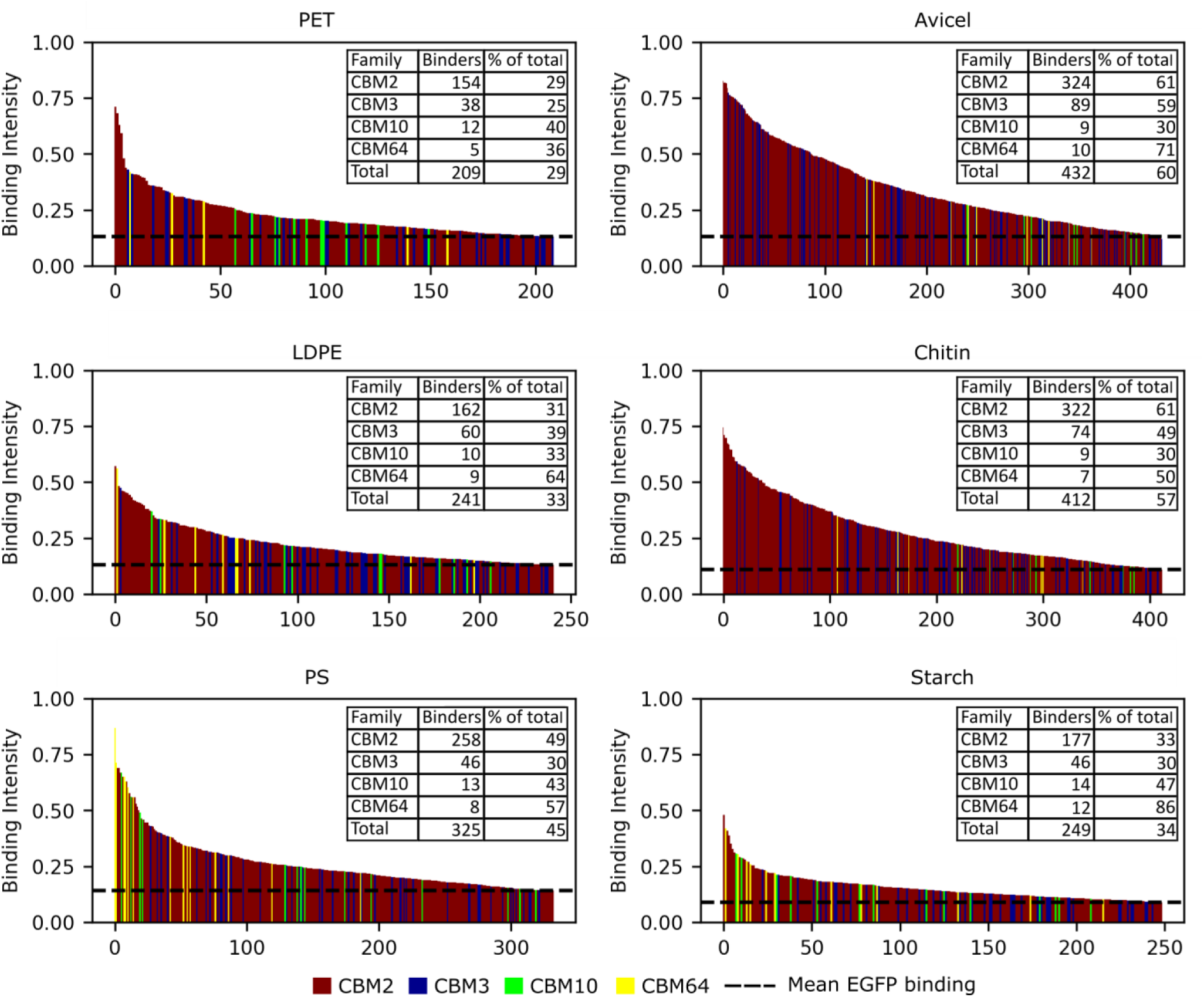
Histograms showing the binding intensities of EGFP-CBM constructs to each of the six substrates analyzed. Binders are identified as having a higher BI than the average of lone EGFP binding over the course of several plates, as represented by the dotted line on each histogram. The number of binders from each family and total binders on each particular substrate are seen in the insert on each plot, along with the proportion of confirmed binders from the total tested in that family. Tables of the full dataset of the proteins binding to each of the substrates can be found in *SI Appendix*, *Dataset S1,* and *Dataset S2*.

A high number of binders were observed for avicel and chitin with 432 and 412 binders, respectively. These polysaccharides also exhibited higher average BI values than the other substrates, consistent with CBM2 and CBM3 being identified as cellulose and chitin binders (38–40). The CBMs that bound most tightly to avicel and chitin were from organisms found in environments rich in these polysaccharides, including soil bacteria, cellulose-degrading sludge communities, and plant pathogens. Both gram-positive and gram-negative bacteria and eukaryotes were represented, with CBMs from marine bacteria among the strongest chitin binders. Previous studies have identified CBMs with comparable affinities for both cellulose and chitin (41, 42), demonstrated here by 270 CBMs binding to both substrates. Over 100 domains were identified as specific binders to either avicel or chitin, mostly within the CBM2 and CBM3 families. However, specific chitin binders all had lower BI values compared to specific avicel binders. Unsurprisingly, significantly fewer starch-binding domains were identified, with 249 in total, many of which bound weakly (Fig. 2). Only 17 CBMs had a BI value on starch over 0.3, with only one approaching 0.5, a CBM2 from the bacteria *Phytohabitans suffuscus*, isolated from orchid roots. However, a much higher proportion of CBM10 and CBM64 bound to starch compared to avicel and chitin, with many of these coming from putative cellulases or xylanases.

The most promising PET binders were found in the CBM2 and CBM3 families (Fig. 2 and *Dataset S1*). The highest BI on PET was 0.71 for a CBM2 from *Streptomyces bingchenggensis*, which also bound strongly to PS and avicel. Another notable protein, a CBM2 from *Rivularia sp.*, had a BI of 0.59 on PET but did not bind to the other two plastics. These BI values are significantly higher than the 0.19 measured for the widely studied PET hydrolase PHL7 (*Dataset S1*). BI values of CBMs binding to LDPE were overall slightly lower than those for PET and PS (Fig. 2 and *Dataset S1*). However, notable examples of high BI values and specific CBMs were identified, including a CBM64 from the bacterium *Fulvivigra maritima*. It demonstrated specific binding to LDPE with a BI of 0.56, with several CBM2 domains also exhibiting specific binding to LDPE.

Among the tested synthetic polymers, the largest number of hits were on PS, with CBM64 domains representing most of the higher BI values (Fig. 2). BI values on PS were generally higher, with 20 proteins having BI values of 0.5 or greater, compared to 5 for PET and 8 for LDPE. The strongest PS binders were a CBM64 from *Spirochaeta thermophila* and a CBM2 from *Micromonosporaceae bacterium* DSM 106523, both displaying negligible binding to PET and LDPE. The CBM10 family had similar number of hits as CBM64 but generally lower BI values (Fig. 2). Notably, three CBM10s bound tightly and specifically to PS (AAA26710, AAO31760, and BAA25188|330-363).

### Specificity of CBMs

To elucidate the structural and phylogenetic relationships of CBMs exhibiting a given binding selectivity, phylogenetic trees of the 727 proteins that bound to at least one of the polymers were constructed. These trees were split into the four CBM families and annotated with HTP holdup assay results (Fig. 3, *SI Appendix,* Fig. S1). When focusing on the natural polymers, the trees generally reflect known trends within CBM families and reveal phylogenetically determined binding subgroupings. In the CBM2 tree, avicel and chitin binding are evenly distributed (Fig. 3), consistent with known preferences in this family (38, 39). Some subgroups show avicel binding dominance, while others exhibited low binding to both avicel and chitin (Fig. 3*A*), suggesting specificity for other polysaccharides like xylan. The CBM3 tree shows a subgroup with pronounced avicel binding (Fig. 3*B*), indicating phylogenetically determined specificity. The CBM10 and CBM64 trees (Fig. 3*C, D*) show stronger and more frequent binding to starch compared to the other two families. The trees displaying the BI to the synthetic polymers also reveal some groupings (*SI Appendix,* Fig. S1), with PS in the CBM2 family showing some strong phylogenetic dependence.

**Fig. 3.**
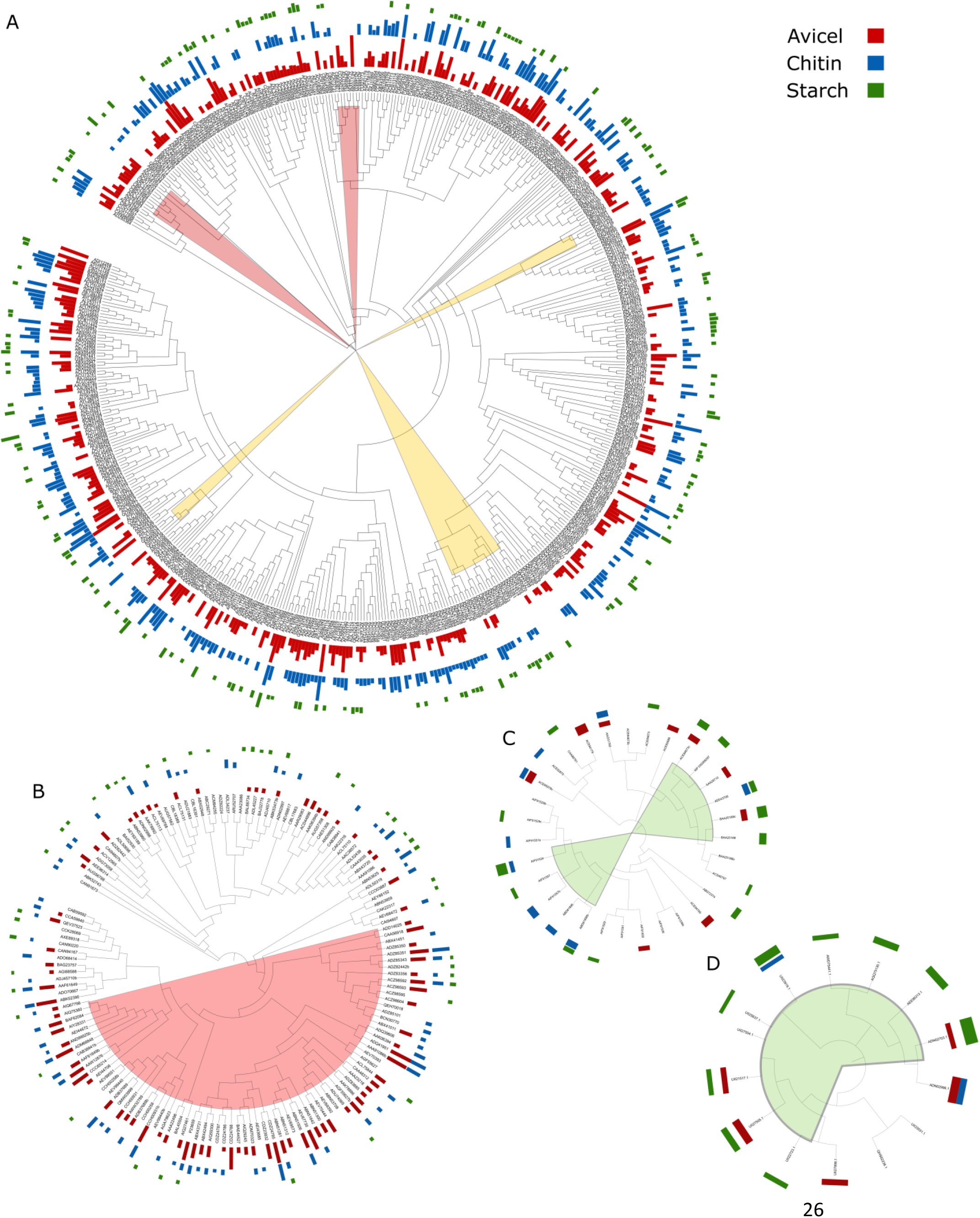
Phylogenetic trees including proteins with at least one confirmed binder in the holdup assays, decorated with the BI of the three polysaccharide substrates, avicel (red), chitin (blue), and starch (green). (*A*) CBM2, (*B*) CBM3, (*C*) CBM10, and (*D*) CBM64. Shading refers to groupings of proteins that bind specifically to the substrate represented by that colour, with the yellow shading representing CBMs that do not bind of any of the substrates. Phylogenetic trees of the same families, decorated with the BI values of the three plastic substrates can be seen in *SI Appendix,* Fig. S6.

Following the observations from the phylogenetic trees, comparisons of Alphafold2 models of selected CBMs were conducted (*SI Appendix,* Fig. S2 to S10) to identify structural specificity determinants, with a focus on the binding surface (22). There were several examples of CBM2 that were missing the canonical aromatic triad (e.g., ADJ46832, ACY9890, and ADJ46833) but showed some substrate preference for chitin and PS, with negligible binding to the other substrates. In particular, polar or charged side chains seem to promote chitin binding in the absence of aromatic residues on the binding surface (*SI Appendix,* Fig. S2 and S3). A significantly modified binding surface is seen in around 10 CBM2, having only a single canonical aromatic residue and a protruding loop of two or three residues containing a tyrosine orthogonal to the plane, with plddt above 95 (*SI Appendix,* Fig. S4). These proteins did not bind any substrates significantly, except for one (AGM06248), which bound tightly to PS and chitin, potentially due to a histidine also protruding orthogonal to the surface plane. An arginine three residues after the first of the canonical binders is seen in around 10 CBM2, serving to rotate the aromatic side chain through 90° (33), which also disrupts binding to PET (*SI Appendix,* Fig. S5).

A majority of CBM3 modules have loop truncations removing one of the aromatic residues from the binding surface, relative to the canonical CBM3 structure (PDB: 1NBC) (e.g., ADZ85343, ABX41011, and ADZ85351). These proteins show consistently strong avicel binding (*SI Appendix,* Fig. S6), in contrast to previous findings (43). The subgroup with the extended loop is underrepresented among both PET and chitin binders. Around 10 of these proteins also contain a short α-helix found just after the binding residue on the fourth β-strand, which protrudes into the binding surface (*SI Appendix,* Fig. S7). This CBM3 subgroup generally showed very weak binding to the polysaccharides, especially when the helix contained a polar residue making up the binding surface. However, two members of the subgroup, which had two hydrophobic residues in the helix contributing to the binding surface, showed moderate avicel binding, while only binding weakly to chitin.

Three of the 17 screened CBM64 members show major structural variation with an extended loop, which adds another tryptophan residue to the binding surface (*SI Appendix,* Fig. S8). Proteins with this third tryptophan did not bind PET or LDPE significantly, but two of them bound tightly to PS. This subgroup also contains one of the modules with the highest BI on starch, with CBM64 binding starch more frequently than the other CBM families.

The electrostatics and hydrophobicity of the CBM surfaces were mapped to determine any associations between these and substrate specificity. Maps displaying the electrostatic potential and Wimley-White hydrophobicity (44) of each of the 1190 CBMs selected for the screen in this study can be found in *Dataset S3*. Each of the CBMs with a BI of 0.30 or more on starch showed an extended hydrophobic surface (*SI Appendix,* Fig. S11), extending away from the canonical binding surface. In terms of the synthetic polymers, some parallels exist between the hydrophobic surfaces of the most tightly binding CBMs on each plastic (*SI Appendix,* Fig. S12). The strongest PS and PET binders seem to have a discontinuous hydrophobic binding surface, with polar patches surrounding it. LDPE binders seem to have a more continuous hydrophobic surface, with CBM2 binding to LDPE possessing a neutral or minimally charged binding surface (*SI Appendix,* Fig. S13).

### Mixed Substrate Assays with CBM Fusion Enzymes

In this study, one CBM from each of the four screened CBM families, with varying specificity to PET or PS, was fused with the commonly studied PET hydrolase, LCC^ICCG^ (45). The activity of these fusion enzymes on crystalline PET powder was assayed and compared to LCC^ICCG^ alone (Fig. 4). Assays were performed at 50 °C, which is below the optimum temperature of LCC^ICCG^, to accommodate the lower thermostability of the binding modules (*SI Appendix,* Fig. S14). To confirm proper folding of the CBMs in the fusion enzymes, Langmuir binding isotherms to avicel were produced (*SI Appendix,* Fig. S15). All enzymes, except LCC^ICCG^-*Fl*CBM64, showed affinity for avicel not seen in LCC^ICCG^ alone. However, *Fl*CBM64 did not bind strongly to avicel in holdup and pulldown assays. Furthermore, Langmuir binding isotherms for each protein on PET and PS were also produced. LCC^ICCG^ was shown to bind non-specifically to PET, PS, and LDPE with varying affinity. The fusion enzymes showed a range of affinities for the two substrates, with LCC^ICCG^-*Rs*CBM2 showing the highest affinity for PET at 7.2 nM, and LCC^ICCG^-*Fl*CBM64 the highest for PS at 8.7 nM.

**Fig. 4.**
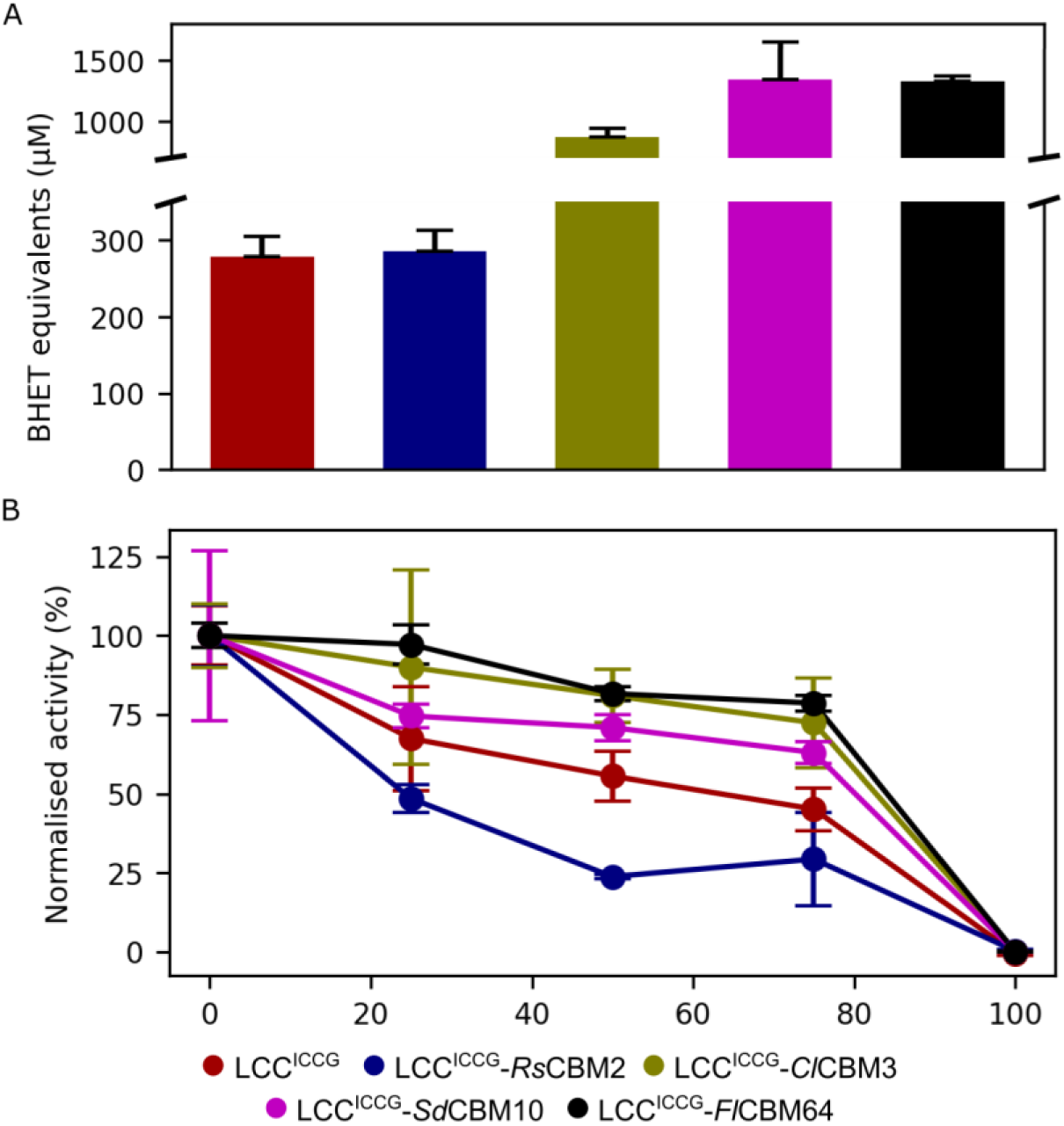
Activity and selectivity of fusion enzymes made with LCC^ICCG^ and CBMs taken from holdup assay. (*A*) Activity of LCC^ICCG^ fusion enzymes on PET. 300 nM of enzymes were incubated in 50 g/L of crystalline PET powder for 1 hour at 50 °C. Each point represents the mean of triplicate experiments. (*B*) Activity of LCC^ICCG^ fusion enzymes on PET in mixtures containing increasing amounts of PS “impurity”. 300 nM of enzymes were incubated in mixtures of PET and PS for 1 hour at 50 °C. The activity of the enzymes was calculated per unit mass of the PET substrate and then normalized to the activity in the experiments without impurity. Each point represents the mean of triplicate experiments.

CBM2s had no effect or were detrimental to activity on crystalline PET powder (Fig. 4*A*), consistent with previous findings (46). However, fusions with other CBMs significantly increased PET degradation, with up to a 5-fold increase for CBM10 and CBM64 fusions (Fig. 4*A*). To mimic a mixed plastic environment, assays were performed with increasing amounts of PS “impurity” replacing PET in the mixture to analyze the specificity of the enzymes (Fig. 4*B*). For the fusion enzymes with *Cl*CBM3 and *Fl*CBM64, no significant reduction in activity was observed upon adding PS impurity up to 75 % of the total reaction mixture, compared to a steady decrease in activity for LCC^ICCG^ alone. Conversely, the fusion enzyme with *Rs*CBM2 showed significantly lower activity on the PET substrate as the PS impurity concentration increased, indicating increased binding to the PS impurity compared to the lone catalytic domain.

## Discussion

### A High Throughput Assay for CBM Interactions

We developed an HTP holdup assay for detecting protein binding to insoluble synthetic polymers and polysaccharides, incorporating multiple controls to ensure data quality and prevent false positives. An in-depth discussion of the design and validation of the assay can be found in the supporting information. This protocol is readily automated on a commercial robot, such as the Tecan Evo200 liquid handling robot used in this study, and can be transferred to most systems or performed manually. Key decisions were made to maintain the HTP nature and applicability to various substrates. To minimize substrate use, substrates in the filter plates were reused and regenerated by washing with 1 M NaCl and 2 M urea. Two variants of a chitin binding domain (ACQ50287) were distributed randomly across the chitin test plate, and the BI was monitored over 10 rounds of washing. No decrease in binding was observed over this process, with only normal variation seen between washings (*SI Appendix,* Fig. S16). In addition, several strongly binding proteins were identified on the other substrates after up to 10 washes, suggesting recycling of these substrates is feasible. However, the washing procedure did cause xylan to form gels within the filter plates making it impossible to recycle. In such cases, or when the availability of substrate is not a factor, fresh loading of new plates for each round of CBMs to be tested could be considered. This study focused on the interaction of type A CBMs with insoluble polysaccharides. However, type B and C CBMs could also be analyzed using soluble glycans immobilized on insoluble beads.

The HTP holdup assay results were validated for 24 representatives using a pulldown assay, with small and acceptable discrepancies observed (see *SI Appendix*, Supplementary information text). BI values for synthetic polymers were generally higher in the holdup assays than in the pulldown assay. This may be due to protein interactions with residual protein on the surface due to recycling in the holdup experiments. The validation confirmed the HTP holdup assay as a reliable method for identifying the binding of CBMs to insoluble polymers. Hence, the data can be used for further analysis of CBM affinity and specificity, and to identify structural features that determine these properties.

Using BI as a measure of CBM binding to substrates was necessary for adapting the holdup assay to insoluble polymers. Previously, BI was used to calculate a dissociation constant (*K*_d_) for protein binding to peptides immobilized on agarose beads (36). However, the heterogeneous nature of the binding sites on the polymeric substrates invalidates assumptions of one-to-one binding, making the calculation of binding site density on the surface of the particles impossible and preventing *K*_d_ calculation from a single-point assay. Though, it would be possible to modify the holdup assay to produce binding curves for determining binding affinity, this would reduce data point production per protein tenfold. For identifying CBMs for enzyme design, detailed calculation of the binding parameters *K*_d_ and *B*_max_ was secondary to analyzing a large number of CBMs, allowing the selection of the binders and ranking the relative ability of each member of a large family of enzymes. The holdup assay could produce up to 10,000 data points per day, allowing BI calculation of approximately 2,500 proteins on one substrate daily, outperforming previous HTP screening techniques that measure binding on a few hundred samples per day at most (47).

### CBMs Demonstrate Substrate Specificity for Polysaccharides and Plastics

The HTP holdup assay identified several hundred CBMs binding to each of the six substrates tested, many of which were previously unidentified. Within the polysaccharides, the preference for starch binding by CBM10 and CBM64 indicates a previously unidentified binding tendency in these families. Grouping of starch binding in the CBM10 and CBM64 trees suggests an evolutionary development of starch binding or a lack of pressure for substrate specificity, resulting in promiscuity. A study on the CBM64 from *Spirochaeta thermophila* (ADN02703.1) showed that this CBM displays properties of both type A and B binding modes, binding to solubilized polysaccharides as well as insoluble flat surfaces (30). This dual binding mode may explain the binding of this and other CBM64 modules to starch, with many amylopectin branches behaving as semi-solubilized polysaccharides on the granule surface (34, 35). The CBMs from each family identified in this study binding to starch all had hydrophobic surfaces extending from the canonical binding surface (*SI Appendix,* Fig. S11), which may facilitate this dual binding as glycans extending from the starch surface wrap around the CBM during binding. Importantly, none of these CBMs originate from starch-active enzymes, so the ability to bind starch could serve as an anchor during degradation of nearby cell material.

Specific binding to chitin by some CBM2 members seems to be driven by polar residues either in place of or surrounding the aromatic binding residues. The presence of polar amide groups on the chitin chain would be more available for hydrogen bonding than the hydroxyl groups in cellulose, which would favor polar interactions in bonding. Within the CBM2 and CBM3 trees, phylogenetic groupings of low chitin and cellulose binding (Fig. 3A and 3B) suggest other substrate specificities within these families, such as xylan. The CAZy database only reports one instance of CBM3 binding to chitin (48); however, around 70 were identified here, with several showing a strong preference over avicel. Many of the CBM10 domains screened are natively found as part of repeat tandems, which do not always have the same specificity or affinity (31), with similar effects seen in this study. Two CBM10 domains of a mannanase from *Vibrio sp.* (BAA25188) were screened alone and in their native conformation as a tandem protein, with only the C-terminal domain showing affinity for starch and avicel.

The presence of phylogenetic groupings in the trees decorated with the BI of synthetic polymers is a surprising result. However, given that substrate specificity is observed with CBMs binding to these substrates, this must be driven by structural and sequence-based features. Most CBM2 modules have the canonical aromatic triad of three tryptophan residues, with most of the strongest binders to each substrate represented in this group. The third position varies, featuring aromatic, polar, or aliphatic side chains. Most CBM2s with glycine in this position show negligible binding to the polysaccharides (*SI Appendix,* Fig. S9), while histidine and serine (*SI Appendix,* Fig. S10) allow strong binding and specificity, indicating a target for future engineering of CBM2.

Within the CBM2 family, a known dimorphism exists between glycine and arginine residues three positions after the first of the three canonical binding aromatic residues, with arginine rotating it through 90° and affecting the planar binding surface (*SI Appendix,* Fig. S5). This rotation has been proposed to reduce binding to cellulose and drive specificity to xylan (33), which was not included in the screen. Of the eleven proteins with arginine at this position, binding to PET and LDPE was significantly reduced, while several bound tightly to PS. Several other indicators of specificity to PS binding were observed, such as an extra aromatic binding residue in CBM64 or a tyrosine protruding from the binding surface in CBM2, with each of these possibly driven by the non-planar conformation of aromatic rings. Each of these is a target for producing CBM variants with a high specific affinity to PS over other plastics.

The hydrophobic and electrostatic maps also suggested further drivers of specificity between the plastics. The strongest PET binders had hydrophobic patches that were more separated compared to the PS binders, with LDPE binders exhibiting a more continuous hydrophobic surface (*SI Appendix,* Fig. S12). Binding specificity is often influenced by the distance between aromatic residues in nature (32). In PET binders, the separated hydrophobic patches could align with the more widely spaced aromatic moieties, as compared to PS, facilitating specificity.

### CBMs Can Be Used to Engineer Specific Plastic Degrading Enzymes

The increased activity upon CBM fusion to a PET hydrolase aligns with previous studies on different PET substrates (24), confirming that HTP screening can identify substrate-binding modules with promising applicability in enzymatic PET degradation. It should be noted that these fusion enzymes may be less applicable than LCC^ICCG^ for degrading PET in a single stream due to their lower thermal stability (*SI Appendix,* Fig. S14) and the previously mentioned effects of high substrate load. However, they could offer advantages in mixed plastic environments, such as municipal solid waste.

The known PET binder *Ba*CBM2, proposed for improving PET hydrolase activity (46), had a BI around the median for CBMs binding PET in this study. This highlights the potential of many CBMs identified here for obtaining high-affinity, high-activity PET hydrolase fusion enzymes. CBMs selectively binding LDPE and PS were also identified. To date, only two enzymes have been fully characterized with activity on PE, and little is known about their binding modes or affinity for their target, leaving scope for future design of fusion enzymes for PE degradation using the CBMs identified in this HTP screen. No enzymes in the PAZy database are currently characterized with activity on PS, but several studies have identified putative enzymes with some PS degradation capability (49). The PS-binding CBMs identified here could aid in designing enzymes for degrading this polymer.

Strong binding of a PET hydrolase to PET does not always correlate with high activity, as dynamic adsorption and desorption are considered more relevant properties (46). Hence, the identification of drivers of CBM affinity and specificity could facilitate the future development of CBMs tailored for engineering enzymes for specific situations. Several fusion enzymes produced in this study had enhanced activity on crystalline PET powder (Fig. 4A), while also maintaining activity in the presence of another plastic, which acts as a competitor to binding (Fig. 4B). Higher activity of LCC^ICCG^-*Cl*CBM3 and LCC^ICCG^-*Fl*CBM64 on PET alone suggests either slower adsorption or faster desorption from the PET surface, likely applying to PS as well. This would allow the enzymes to maintain higher activity on PET despite transient inactivity while bound to PS, due to increased dynamic interactions with both substrates. Development of these enzymes would be highly valuable for the enzymatic degradation of mixed plastic wastes. Furthermore, these insights could guide future engineering efforts to develop enzymes active on PS and LDPE, substrates for which enzyme discovery is ongoing.

## Materials and Methods

### HTP Expression and Purification of Proteins

Expression and purification of type A CBMs fused with EGFP (see *SI Appendix* for the description of the selection process) were done using a modified version of the method previously reported (50). All cultures were grown in LB media supplemented with 50 µg/mL of kanamycin. Initially, agar stabs were used to inoculate 1 mL pre-cultures in 96-deep-well (DW) plates, which were covered with a breathable film and grown overnight at 37 °C, 200 rpm. Glycerol stocks were then made from each culture and used for subsequent pre-cultures. 200 µL of each pre-culture were used to inoculate 2 mL cultures in 24-DW plates, which were grown at 37 °C, 160 rpm for 2 hours before being induced with 1 µM of IPTG and grown overnight at 20 °C, 160 rpm. Cells were harvested by centrifugation at 5000 rpm, and resuspended in 1 mL of lysis buffer (50 mM Tris-HCl, 500 mM NaCl, 0.1 mM phenylmethylsulfonyl fluoride (PMSF), 0.25 mg/mL lysozyme, pH 8.0), then frozen at −20 °C. DNase I and MgSO_4_ were added to the frozen pellets at final concentrations of 20 µg/mL and 20 mM, respectively, and the cells were lysed by thawing at 37 °C for 10 minutes. Next, 200 µL of 25 % (dry bead volume/slurry volume) nickel chelating sepharose fast flow agarose bead slurry (Cytiva, USA) was added to the lysates, and the protein was allowed to bind with gentle shaking at 20 °C for 10 minutes. The lysates were then transferred to a 96-well filter plate (Machery Nagel, Germany) in a Multiscreen vacuum manifold (Sigma-Aldrich, Germany) fitted with a DW adaptor. Vacuum was applied, and the flow-through was collected in a 96-DW plate. The Ni resin beads were washed sequentially with 10 mM imidazole and 50 mM imidazole in HTP binding buffer (50 mM Tris-HCl, 500 mM NaCl, pH 8.0), and the protein was eluted in 500 µL fractions using HTP binding buffer with 500 mM imidazole. The purified protein was then stored at −20 °C prior to analysis.

### HTP Screening of CBMs against Plastics and Polysaccharides

All liquid handling for the holdup assays was performed using an Evo200 liquid handling robot (Tecan, Switzerland) with a 96-tip pipetting head. Substrates were suspended at a concentration of 13 mg/mL in either assay buffer (50 mM Tris-HCl, 500 mM NaCl, pH 8.0) or ethanol, depending on the substrate density to ensure even suspension. The settled substrate in the transfer liquid was resuspended by vigorous pipetting before transferring 75 µL to a 384-well MZHV Millipore plate with a low-binding membrane (0.45 μm Durapore PVDF membrane, Millipore, USA), delivering approximately 1 mg to each well. The 384-well plates were divided into 96 groups of 4 wells in a square, with the bottom right well not containing any substrate, serving as a control (Fig. 1). After substrate addition, the 384-well plates were placed on a vacuum manifold and aspirated to remove the transfer liquid. Each plate was then sequentially washed with 100 µL 1 M NaCl, 2 M urea, and assay buffer, with removal of liquid on the Tecan vacuum manifold, followed by a final wash with assay buffer and drying by centrifugation at 800 g for 5 minutes. Purified protein, spiked with 100 nM of an in-house produced mCherry (Uniprot: D1MPT3), was added to the 384-well filter plate, and the plate was shaken at 1200 rpm at room temperature for 15 minutes. The filter plate was then centrifuged at 800 g for 5 minutes with a 384-well receiver plate attached to the bottom. This receiver plate was then read on a PHERAstar FSX Microplate Reader (BMG Labtech) at two excitation/emission pairings: 575 nm/620 nm and 485 nm/520 nm to measure mCherry and EGFP fluorescence, respectively. After each run, the washing procedure of sequential urea, NaCl, and assay buffer washing was repeated, followed by a final wash with assay buffer and drying by centrifugation. The regenerated plate was then reused for the next cohort of EGFP-CBM constructs to be tested.

Given that only single-point concentration determinations were made during the HU assays, the *K*_d_ of the CBMs binding to each substrate could not be determined. Instead, the related parameter of binding intensity (BI) was used. This is defined as:

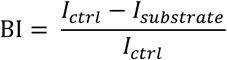

Where *I_ctrl_* and *I_substrate_* refer to the intensity of the EGFP fluorescent signal in the control wells without substrate and the substrate-containing wells, respectively. The EGFP signal intensities are normalized to the median mCherry signal in a group of four wells to account for small differences in pipette volume from the 96-tip pipetting head or incomplete transfer of the liquid from the filter plate to the receiver plate during centrifugation. The BI represents the fraction of the EGFP-CBM construct bound to the substrate in the wells, allowing comparison of the relative binding strengths of the proteins at the 0.5 µM concentration used. The mean BI from each of the three wells containing substrate was reported for that particular protein on that substrate. Error bars representing the variance between the points retained after data curation are shown in the raw data and results files in *Dataset S2*.

Several controls were used to ensure data quality during the HU assays. mCherry was added to every well at the same concentration, allowing for the correction of volume discrepancies by comparing its fluorescence intensity (575/620 nm band pass filter) to the median fluorescence intensity for that group. Signals not within 80 % to 120 % of the median value of the four wells were discarded and not used to determine the binding intensity. If one of the three test wells was discarded from the data analysis, either the mean of two values or a single replicate was used as the reported BI. Purified EGFP protein was randomly distributed in the 96-well plates used to fill the 384-well filter plates, allowing for the determination of EGFP binding to each substrate and serving as the threshold above which a CBM was considered a binder to a particular substrate. Additionally, four known PET hydrolase enzymes, LCC^ICCG^, *Is*PETase, PHL7 and *Tf*Cut2 (12, 45, 51, 52), were analyzed for their BI on each of the substrates as controls.

### Structural Analysis of CBMs

The structure of each of the CBMs expressed in the study was modeled using AlphaFold2 (53). These models were structurally aligned using the sequence-independent and structure-based *super* algorithm in Pymol2, with 5 cycles, each having a 2.0 Å refinement cut-off for identified atom pairs. This structural alignment allowed easy comparison of 2D projections of the surface features (44, 54–57) of the CBM models, including electrostatic potential and hydrophobicity, which were visualized using the SURFMAP software (58). The electrostatic potential maps were visualized through calculation of the potential at pH 7.5. The individual 2D projections of the CBM surfaces were inspected manually.

### Benchmarking HTP Binding Results using Pulldown Assay

To benchmark the results obtained in the holdup assay, 24 of the EGFP-CBM proteins analyzed in the holdup assay were also tested using a pulldown assay. The proteins were produced and purified as described in Section 2.7. 4 mg of each substrate in 100 µL of HTP binding buffer were added to 96-well low-binding microtiter plates, along with 100 µL of EGFP-CBM fusion protein, to a final protein concentration of 0.5 µM. Control wells without protein were also included. The plate was shaken for 15 minutes at 25 °C and 300 rpm in a thermomixer (Eppendorf, Germany), and then spun at 1000 rpm for 5 minutes in a centrifuge. Next, 150 µL of the supernatant was transferred to a new 96-well low-binding microtiter plate and centrifuged again. Then, 100 µL was transferred to a microtiter plate suitable for reading in a spectrofluorimeter. The concentration of unbound protein was determined by fluorescence using an FP-8500 Spectrofluorimeter equipped with an FMP-825 plate reader (JASCO Corporation, Japan) with excitation at 450 nm and emission at 510 nm. BI was calculated in the same way as in the holdup assay.

### Langmuir Isotherms

The affinity of the proteins used in the mixed substrate assays for PET was determined by a pulldown assay. A dilution series between 10 nM and maximum 1000 nM of each protein in assay buffer was mixed with 50 mg of PET^CP^ in 300 µL volume in a non-binding microtiter plate (In Vitro, Denmark). The samples were incubated for 1 hour at 50 °C in a thermomixer (Eppendorf, Germany), and then spun at 3000 g for 5 minutes and the supernatant moved to a new plate. The amount of unbound protein was determined using a BCA assay kit (Thermofisher, USA), specifically mixing working reagent and supernatant in a 1:1 ratio followed by incubation at 37 °C for 2 hours before reading absorbance at 562 nm in a spectrophotometer (Epoch Biotek, USA). The amount of bound protein in each well was calculated by subtracting the unbound from the total protein added, and the dissociation constant (*K*_d_) was determined by fitting eq. 3 to the data.

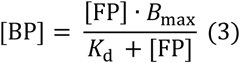

Where [BP] is the coverage of protein on the substrate (nmol/g), [FP] is the concentration of unbound protein (nM) and *B_max_* is the amount of proteins required to fully saturate the surface of the substrate. For the isotherms, the concentration of bound protein was expressed as nmol/g. A minimum of seven points were used to produce each binding isotherm for *K*_d_ and *B_max_* determinations, and assays were run in triplicate.

### PET Degradation in a Mixed Substrate Assay

Four CBMs, which demonstrated strong binding and selectivity in the holdup assay, were used to produce fusion enzymes with the PET-degrading enzyme LCC^ICCG^ (9, 10). Additionally, a fusion with the known PET binder *Ba*CBM2 was produced as a control. Details of the LCC^ICCG^ fusion enzymes are provided in *SI Appendix,* Table S2. They were produced and purified as described in the supplemental materials and methods. The activity of these enzymes, along with LCC^ICCG^ alone, on amorphous PET powder was assayed with increasing amounts of PS or avicel added as “impurities” to the reaction mixture. Assays were performed in 96-well low-binding microtiter plates in a 200 µL volume with an enzyme concentration of 300 nM. A total solid mass of 50 mg was maintained, with PS or avicel replacing PET at 25 %, 50 %, 75 %, and 100 % of impurity. The microtiter plates were shaken at 1000 rpm and 50 °C for 1 hour, then centrifuged at 1000 rpm for 5 minutes. Next 150 µL of the supernatant was transferred to a new 96-well low-binding microtiter plate and centrifuged again. Then, 100 µL was transferred to a new UV microtiter plate. The absorbance of degradation products was measured at 240 nm using a spectrophotometer (Epoch Biotek, USA), and the concentration was calculated using a standard curve. Activities were calculated and normalized to the mass of PET substrate under the specific assay conditions. These activities were then normalized to the activity of the enzyme on PET alone and plotted against the percentage of impurity in the mixture.

Additional Materials and Methods can also be found in the Supporting Information.

## Acknowledgments

We would like to thank Professor Bernard Henrissat for his invaluable help in providing sequences of the CBMs, his expert advice on the bioinformatic approaches, and his valuable feedback on the manuscript.

We would also like to thank Bo Pilgaard for his help in preparing the AlphaFold2 models of the CBMs used in this study.

We would like to thank laboratory technician trainee Sabrina Rostved for her help in the expression and purification of proteins for the pulldown assays.

This work was supported by the Danish Independent Research Council [grant number: 1032-00273B]. Furthermore, the gene synthesis work (proposal:10.46936/10.25585/60008756) conducted by the U.S. Department of Energy Joint Genome Institute (https://ror.org/04xm1d337), a DOE Office of Science User Facility, was supported by the Office of Science of the U.S. Department of Energy operated under Contract No. DE-AC02-05CH11231. This publication has received funding from France 2030, the French Government program managed by the French National Research Agency (ANR-16-CONV-0001) and from Excellence Initiative of Aix-Marseille University - A*MIDEX. J.F.M was funded by Consejo Nacional de Humanidades Ciencias y Tecnologias (CONAHCYT) Becas al Extranjero Convenios GOBIERNO FRANCES 2021 - 1 grant 795494. Finally, we would like to thank the French Infrastructure for Integrated Structural Biology (FRISBI) (Grant ANR-10-INSB-05-01) for funding of the equipment at AFMB.

## Disclaimer notice

This document contains results from work performed under the auspices of the U.S. Department of Energy’s Office of Science, Biological and Environmental Research Program and by the University of California, Lawrence Berkeley National Laboratory, Lawrence Livermore National Laboratory and Los Alamos National Laboratory. Neither CONTRACTOR, DOE, the U.S. Government, nor any person acting on their behalf: (a) make any warranty or representation, express or implied, with respect to the information contained in this document; or (b) assume any liabilities with respect to the use of, or damages resulting from the use of any information contained in the document.

## Data, Materials, and Software Availability

All study data are included in the article or *SI Appendix* and *Datasets S1 – S4*.

Links to the datasets can be found in the Dataset legends in the *SI Appendix*.

