## Supplementary Information for "A high throughput investigation of the binding specificity of carbohydrate-binding modules for synthetic and natural polymers"

**This PDF file includes:**

Supporting text

Figures S1 to S18

Tables S1 to S2

Legends for Datasets S1 to S4

Supplemental Materials and Methods

SI References

**Other supporting materials for this manuscript include the following:**

**Dataset S1.** List of carbohydrate-binding modules (CBMs) included in the study.

**Dataset S2.** Raw data from holdup assays.

**Dataset S3.** SURFMAP results of electrostatic potential and hydrophobicity for each CBM.

**Dataset S4.** Caliper data for CBM expression analysis

### Supporting Information Text

**Selection of CBMs for HTP screening.** Type A CBMs are expected to have the most efficacy in binding to insoluble hydrophobic polymers due to their planar binding surfaces (1). Therefore, four families representing Type A CBMs – CBM2, CBM3, CBM10, and CBM64 – were chosen for screening. Single members of these families have already been shown to have a $K_d$ for PET in the nM range (2), indicating their potential as targets for other plastics. These families also exhibit diverse substrate specificities for polysaccharides (3–5), which are abundant in the environment and could play a role for *in situ* plastic waste degradation. CBM2 and CBM3 families are relatively large, with over 14,000 and 3,000 members, respectively. Therefore, for these families and CBM64, targets to screen were selected by filtering sequences for redundancy at a 70 % identity threshold using CD-HIT software (6). For the CBM10 family, representative sequences from each branch of a phylogenetic tree were chosen (3). Since CBM10 modules are often found in tandem, both the native dyad and triad of modules and individual modules were assayed. In total, 1,210 proteins were selected for expression and analysis by the holdup assay, including 909 CBM2, 214 CBM3, 68 CBM10, 18 CBM64 and 20 CBM fusions with common PET hydrolases. The synthesis at Joint Genome Institute (JGI) had a success rate of 90.7 %, which then resulted in 1096 constructs for use in the study. The full list of designed and synthesized proteins can be found in *Dataset S1*.

**HTP CBM Expression.** The process of production of 1096 EGFP-CBM complexes, from agar stabs to purified protein, was completed in 5 days. The concentration of fluorescent protein in each eluted fraction was determined, ranging from undetectable to 41 µM. Concentrations of each eluted protein are detailed in *Dataset S1*. In total 797 proteins were produced with sufficient concentration and checked for purity and integrity (% of EGFP-CBM versus EGFP alone) for use in the holdup assay, see *Dataset S4*. It should be stated that protein purification is not necessarily required given the use of the EGFP label. However, to minimize the amount of proteolytic degradation, and to reduce any non-specific binding of *E. coli* proteins to the substrates, purification was done in this study.

**Design of HTP Holdup Assay.** Inspired by a HTP holdup assay developed for identifying protein binding to peptides immobilized on insoluble beads (7), a holdup assay to screen for proteins binding to insoluble polymers was developed. The protocol is almost fully automated on a tecan workstation, using a modification of the program described previously (8). Due to the complexity of protein-polymer interactions and the impossibility of determining the total number of protein binding sites on each polymer fragment, several precautions were taken, and controls were conducted to validate the method.

To ensure accurate pipetting of the suspended substrate into the 384-well filter plates, the mass of the remaining substrate in the suspensions after the plates had been filled was checked by drying the suspension at 105 °C overnight. The concentration remained consistent before and after dispensing, confirming the pipetting accuracy. Visual inspection confirmed the correct filling of wells, although consistency in substrate amounts per well could not be verified. Furthermore, the red fluorescent signal from mCherry, doped into each purified EGFP-CBM protein, served as a control to ensure consistent pipetting of protein and full transfer of the liquid from the holdup assay into the receiver plate during centrifugation (Fig. 1). False positives could occur if test wells have a lower volume than the control well in a group of four. Therefore, wells with a mCherry fluorescence outside a range of 80 % to 120 % of the median were excluded from the mean BI calculation, with the entire group of four wells being removed if the control well without substrate was outside this range. Finally, a large excess of substrate relative to protein minimized the impact of discrepancies on binding: over-filling had no effect, and under-filling would only lead to tolerable false negatives in this HTP study.

Given that many proteins, including the fusion partner EGFP, bind non-specifically to synthetic polymers (9), lone EGFP modules were randomly distributed across plates to determine the mean BI of the reporter molecule to the reused substrates. EGFP showed low binding to each substrate, with PS having the highest BI of 0.14 and starch the lowest at 0.09. EGFP BI values on each substrate were used as thresholds for when a CBM was considered a binder in the holdup assay (*SI Appendix,* Table S2).

**Comparisons of HTP Holdup Results with Pulldown results.** The HTP holdup assay results were compared with a pulldown assay, which is typically used to determine protein affinity for insoluble substrates. 24 EGFP-CBM fusion proteins, selected from confirmed binders in the HTP holdup assay to cover a range of BI values and substrate specificities, were produced on a larger scale and evaluated by the pulldown assay. SDS-PAGE analyses of these expressions and purifications can be seen in *SI Appendix,* Fig. S18. Of the three natural substrates, only avicel and chitin were analyzed by pulldown, as the holdup results for starch were generally too low for pulldown analysis. LDPE was excluded due to its insufficient density for proper sedimentation by centrifugation. A heatmap of the results is shown in *SI Appendix,* Fig. S17. The heatmaps indicate general agreement between the assays in determining BI values. No binders identified by the holdup assay were non-binders in the pulldown assay, indicating a low likelihood of false positives in the holdup data.

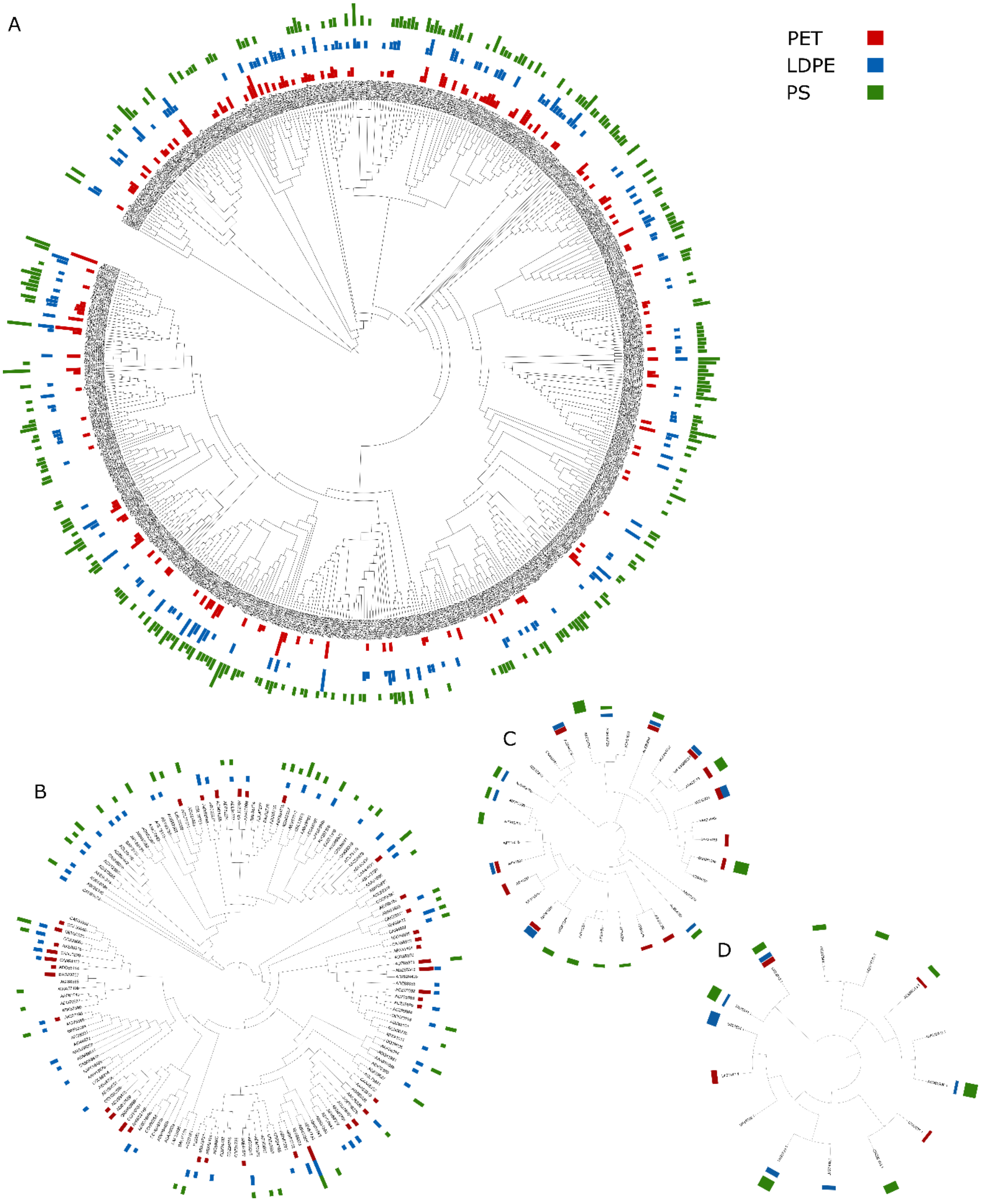
A
B
C
D
PET
LDPE
PS

**Fig. S1.** Phylogenetic trees including proteins with at least one confirmed binder in the holdup assays, decorated with the BI of the three plastic substrates, PET (Red), LDPE (Blue) and PS (Green). (A) CBM2, (B) CBM3, (C) CBM10, (D) CBM64.

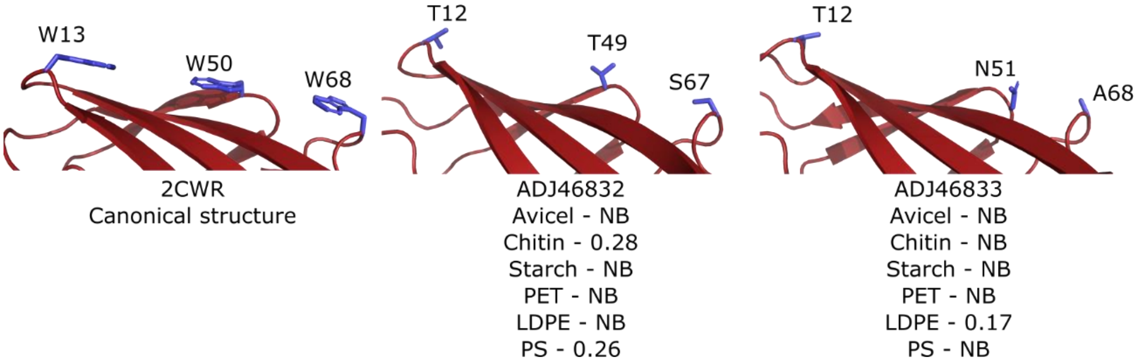


**Fig. S2.** Examples of CBM2 proteins without the canonical aromatic triad, with average BI values on each of the six substrates below, a BI below the biding threshold is represented by NB.

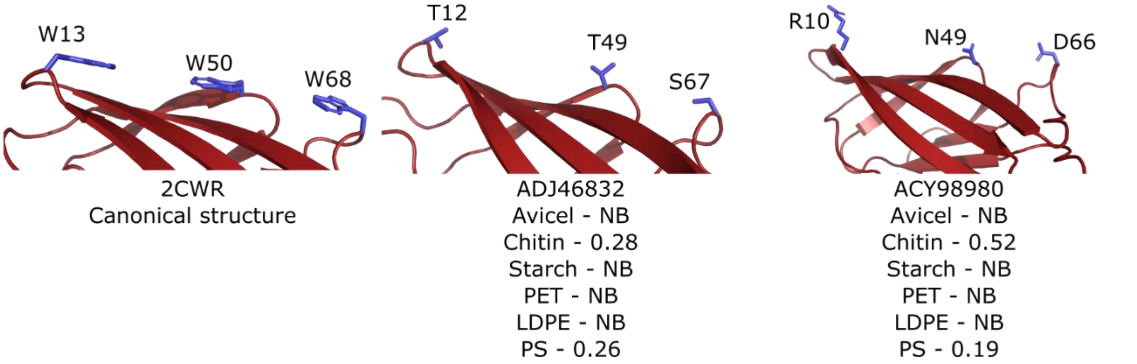


**Fig. S3.** CBM2 proteins that bind specifically to chitin, highlighting the polar residues on the binding surface in place of aromatics, with average BI values on each of the six substrates below, a BI below the biding threshold is represented by NB.

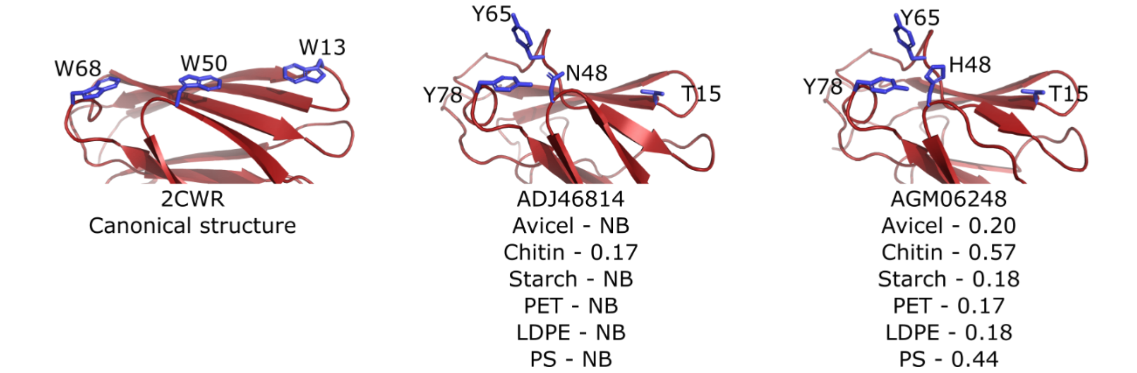


**Fig. S4.** Examples of CBM2 proteins with a modified binding surface, which includes a loop containing a tyrosine protruding from the surface, average BI values on each of the six substrates below, a BI below the biding threshold is represented by NB.

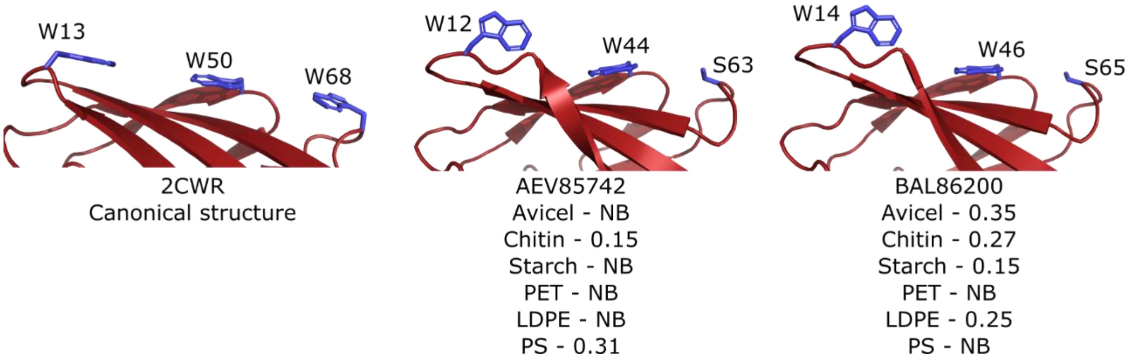


**Fig. S5.** Examples of CBM2 proteins with an arginine three residues after the first binding site, rotating the tryptophan through 90 °, with average BI values on each of the six substrates below, a BI below the biding threshold is represented by NB.

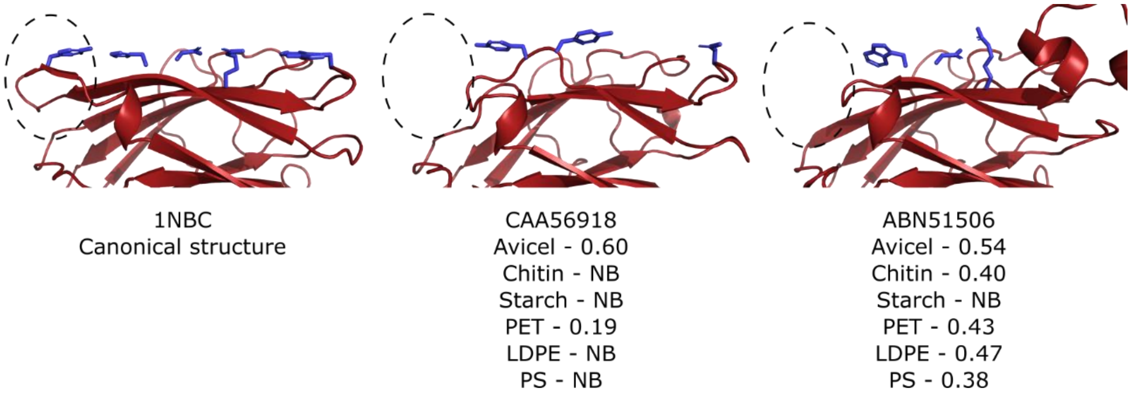


**Fig. S6.** Examples of CBM3 proteins with a truncated loop, with average BI values on each of the six substrates below, a BI below the biding threshold is represented by NB.

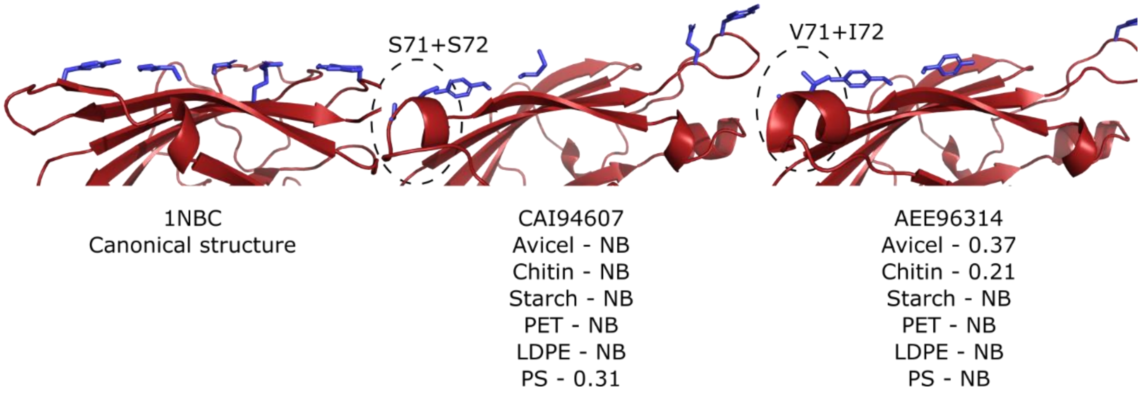


**Fig. S7.** Examples of CBM3 proteins with a modified binding surface, which includes a helix containing polar or aliphatic residues protruding from the surface. Average BI values on each of the six substrates are shown below, a BI below the biding threshold is represented by NB.

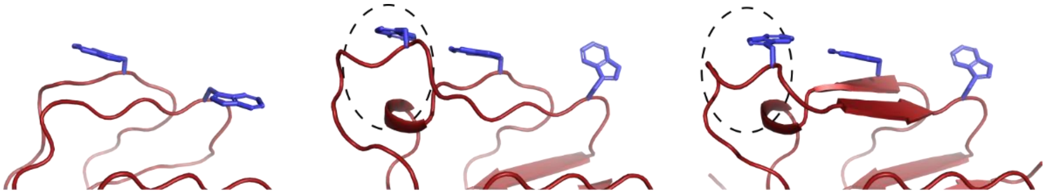

| QHS62238 | AEE96313 | ADN02996 |
|---|---|---|
| Avicel - NB | Avicel - NB | Avicel - 0.28 |
| Chitin - NB | Chitin - NB | Chitin - 0.22 |
| Starch - NB | Starch - 0.26 | Starch - NB |
| PET - NB | PET - 0.15 | PET - NB |
| LDPE - NB | LDPE - NB | LDPE - 0.17 |
| PS - 0.49 | PS - 0.26 | PS - 0.71 |

**Fig. S8.** Examples of CBM64 proteins with an extended loop, conferring an extra aromatic binding residue onto the surface. With average BI values on each of the six substrates below, a BI below the biding threshold is represented by NB.

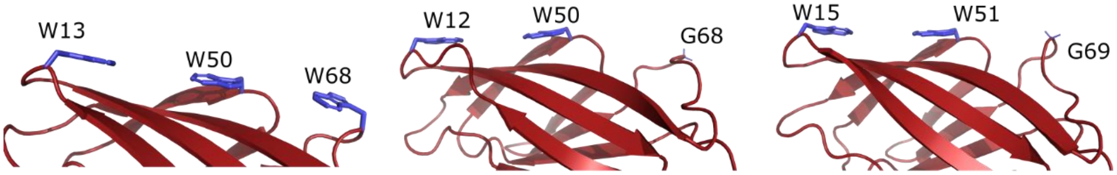


| 2CWR | CAN95423 | ACU35994 |
|---|---|---|
| Canonical structure | Avicel - NB | Avicel - NB |
| | Chitin - NB | Chitin - 0.17 |
| | Starch - NB | Starch - NB |
| | PET - NB | PET - 0.14 |
| | LDPE - NB | LDPE - NB |
| | PS - 0.32 | PS - NB |

**Fig. S9.** Examples of CBM2 proteins with a glycine at the third of the binding sites, average BI values on each of the six substrates below, a BI below the biding threshold is represented by NB.

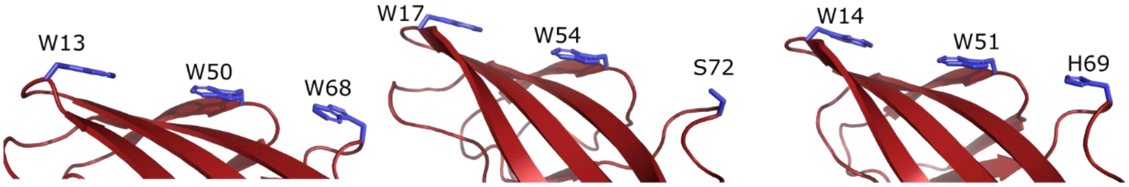


| 2CWR | CAN99125 | AGM06730 |
|---|---|---|
| Canonical structure | Avicel - NB | Avicel - 0.53 |
| | Chitin - NB | Chitin - 0.38 |
| | Starch - NB | Starch - 0.09 |
| | PET - NB | PET - 0.28 |
| | LDPE - NB | LDPE - 0.25 |
| | PS - 0.36 | PS - 0.16 |

**Fig. S10.** CBM2 proteins binding specifically to PS and tightly to avicel and chitin, average BI values on each of the six substrates below, a BI below the biding threshold is represented by NB.

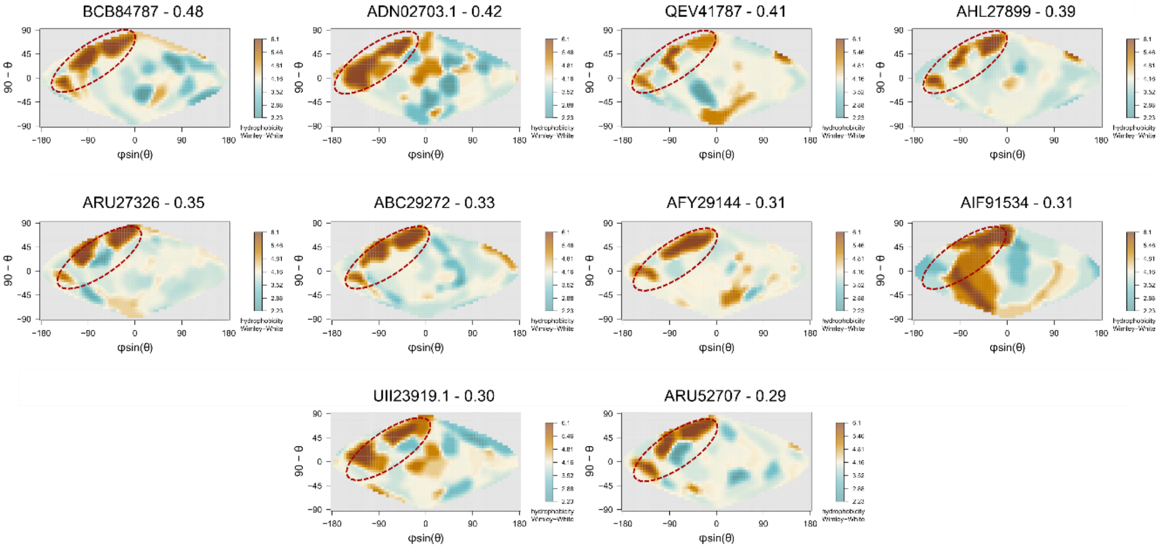


**Fig. S11.** Sinusoidal projections of the surface of proteins binding to starch, with Wimley-White hydrophobicity (10) mapped onto them using the SURFMAP software (11). The Accession number of each, and the measured BI on starch are above each map, with the canonical binding surface highlighted by the dotted red circle on each.

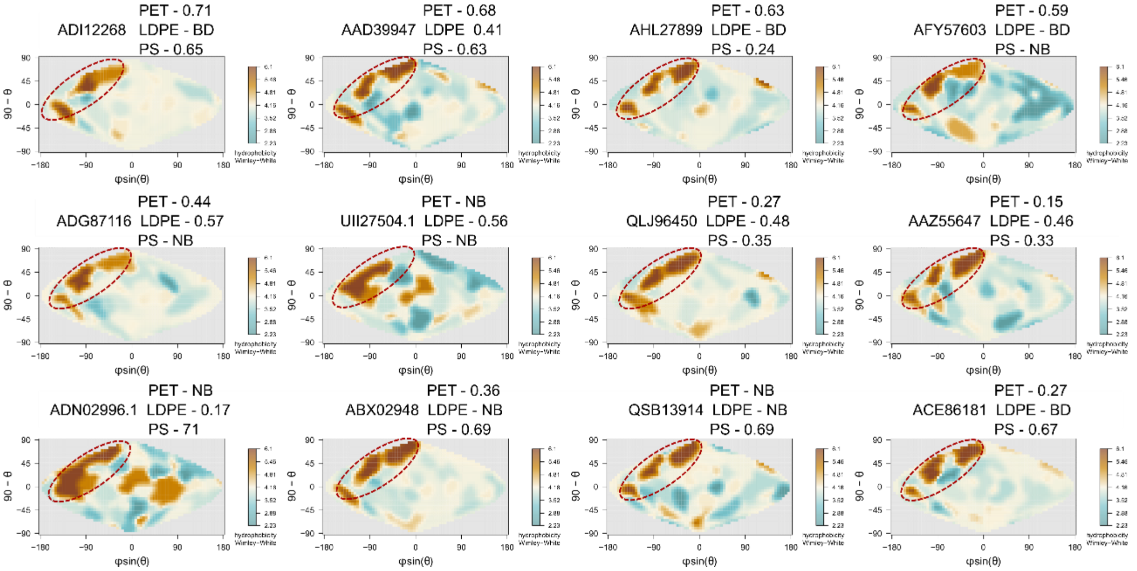


**Fig. S12.** Sinusoidal projections of the surface of proteins with the highest binding intensities to the three plastics (PET binders - top row, LDPE binders – middle row, PS binders – bottom row) with Wimley-White hydrophobicity (10) mapped onto them using the SURFMAP software (11). The Accession number of each, and the measured BI on the three plastics are above each map, with the canonical binding surface highlighted by the dotted red circle on each.

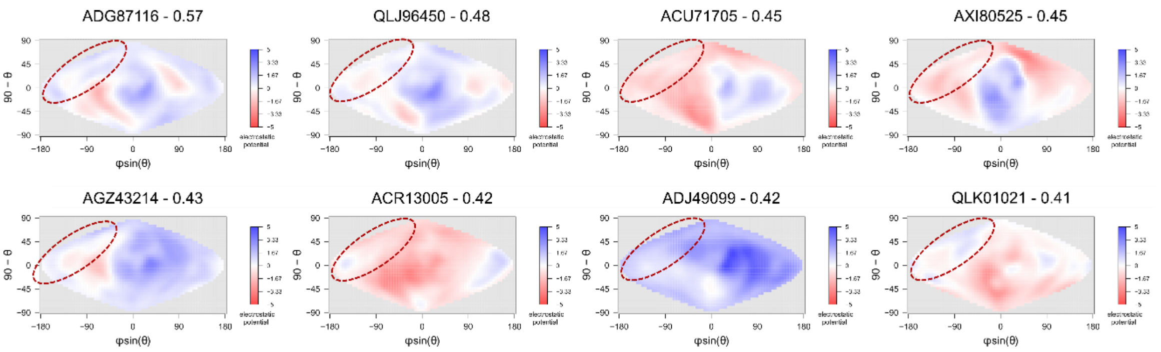


**Fig. S13.** Sinusoidal projections of the surface of proteins binding to starch, with electrostatic potential mapped onto them using the SURFMAP software (11). The Accession number of each, and the measured BI on LDPE are above each map, with the canonical binding surface highlighted by the dotted red circle on each.

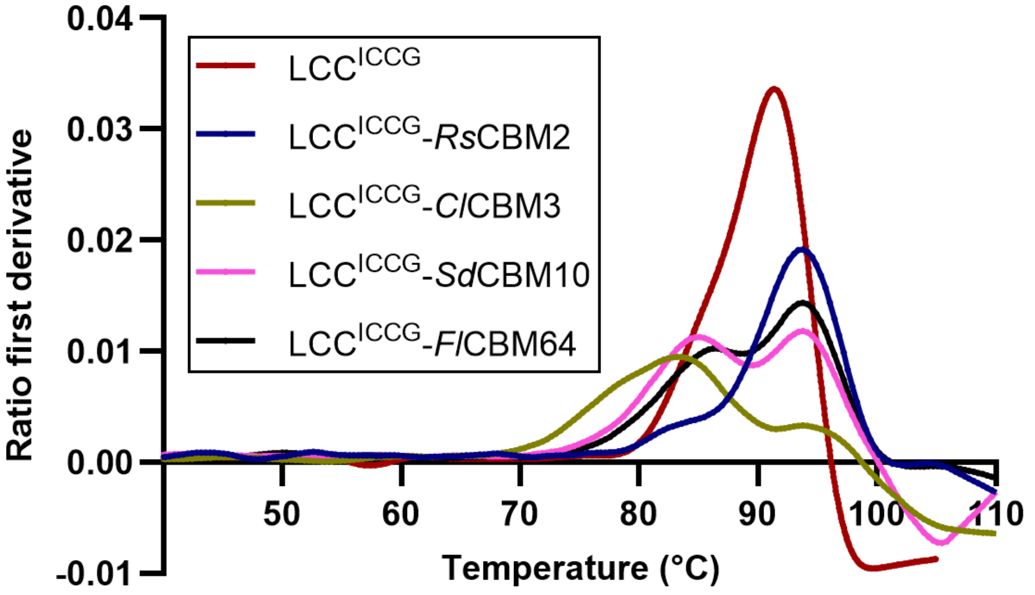


**Fig. S14.** DSF unfolding curves of each of the four fusion proteins produced in this study. Curves are averages of two replicates. Peaks represent the $T_m$ of the different modules within the fusion proteins, with the LCC$^{ICCG}$ module displaying the $T_m$ of approximately 90 °C.

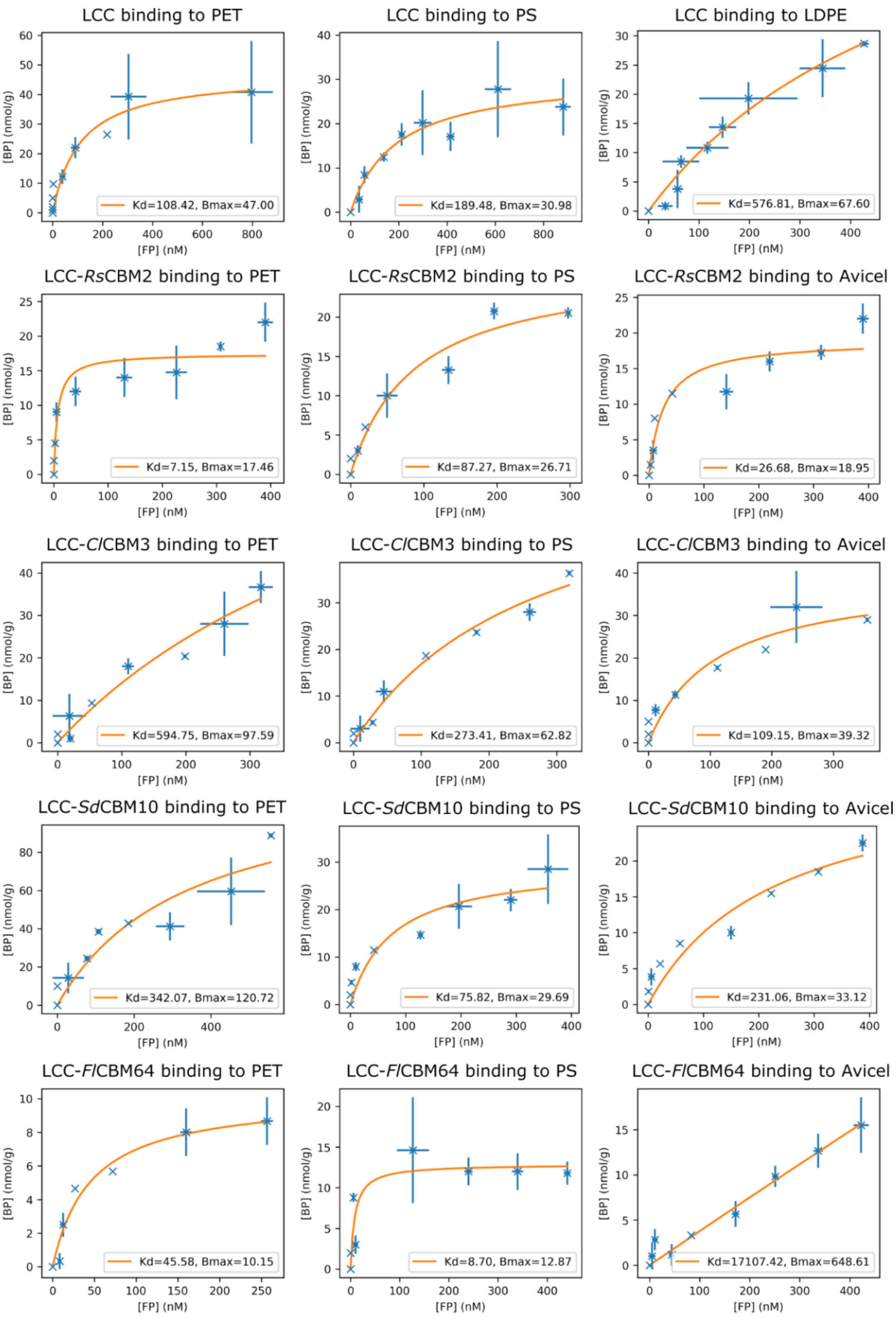
LCC binding to PET
Kd=108.42, Bmax=47.00
LCC binding to PS
Kd=189.48, Bmax=30.98
LCC binding to LDPE
Kd=576.81, Bmax=67.60
LCC-RsCBM2 binding to PET
Kd=7.15, Bmax=17.46
LCC-RsCBM2 binding to PS
Kd=87.27, Bmax=26.71
LCC-RsCBM2 binding to Avicel
Kd=26.68, Bmax=18.95
LCC-ClCBM3 binding to PET
Kd=594.75, Bmax=97.59
LCC-ClCBM3 binding to PS
Kd=273.41, Bmax=62.82
LCC-ClCBM3 binding to Avicel
Kd=109.15, Bmax=39.32
LCC-SdCBM10 binding to PET
Kd=342.07, Bmax=120.72
LCC-SdCBM10 binding to PS
Kd=75.82, Bmax=29.69
LCC-SdCBM10 binding to Avicel
Kd=231.06, Bmax=33.12
LCC-FlCBM64 binding to PET
Kd=45.58, Bmax=10.15
LCC-FlCBM64 binding to PS
Kd=8.70, Bmax=12.87
LCC-FlCBM64 binding to Avicel
Kd=17107.42, Bmax=648.61
[BP] (nmol/g)
[FP] (nM)

**Fig. S15.** Langmuir isotherms of LCC$^{ICCG}$ and the four fusion proteins produced in this study binding to PET, PS and avicel. LCC$^{ICCG}$ had no affinity for avicel. Each point is an average of at least two replicates, with error bars representing the standard deviation.

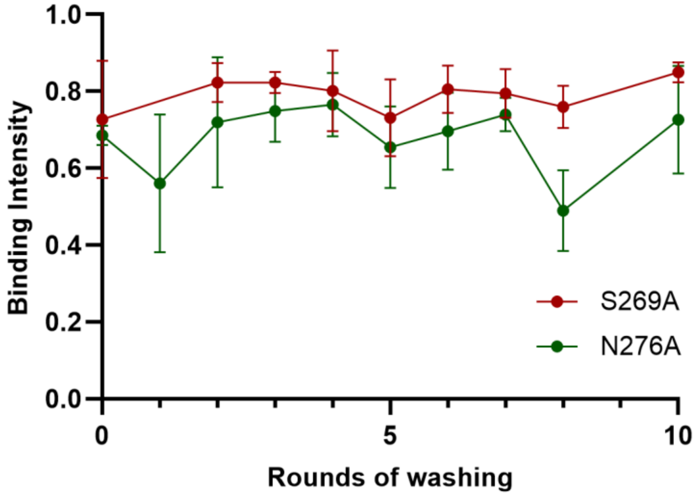


**Fig. S16.** Analysis of the effect of the washing procedure on the substrates. Two variants of a chitin binding CBM (ACQ50287) were randomly distributed over the 384-well MZHV Millipore low binding filter plates, and the binding intensity measured after successive washing cycles of chitin.

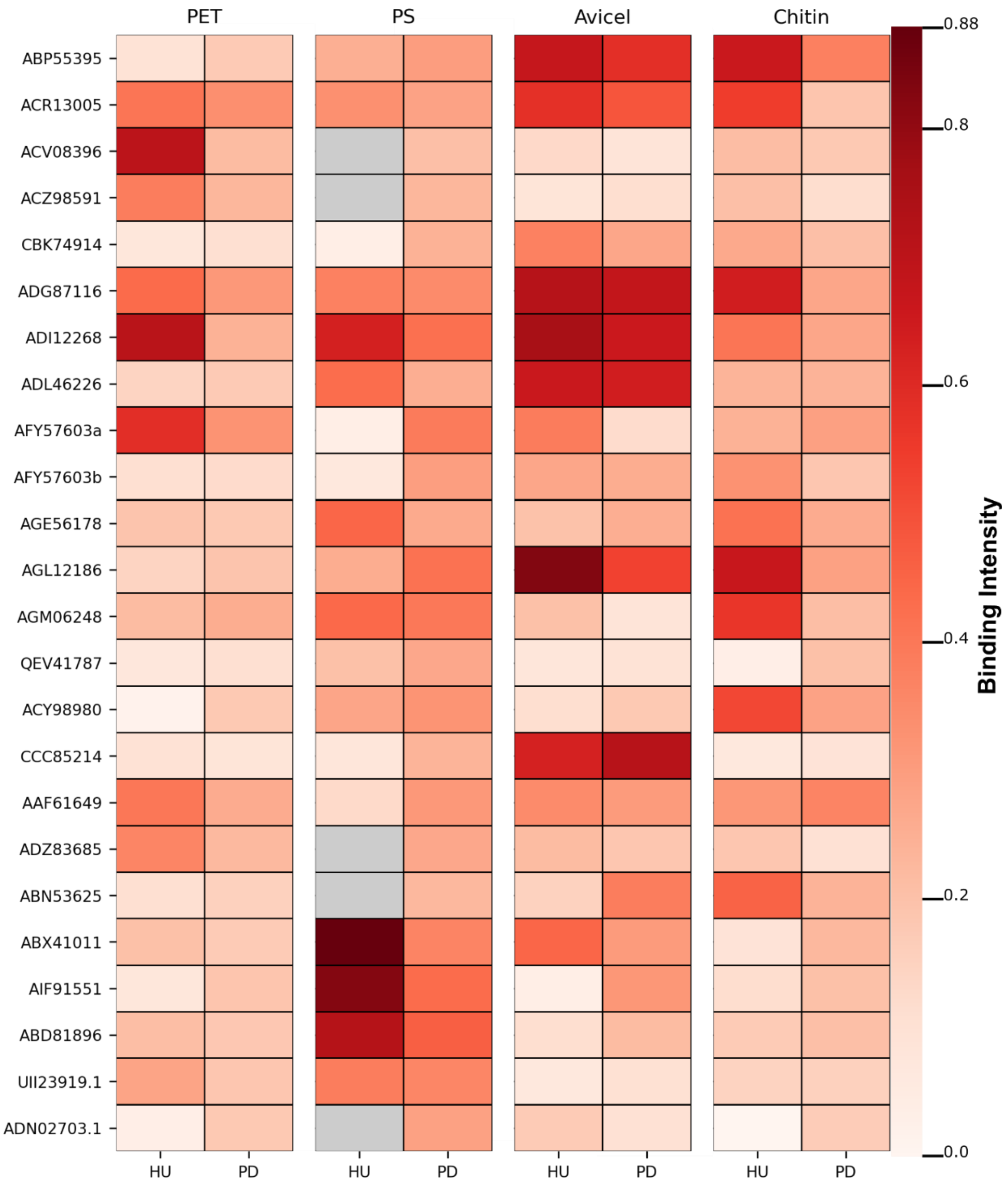


**Fig. S17.** Comparison of the BI as measured by holdup (HU) and pulldown (PD) assays. A heatmap showing the BI of the 24 proteins selected for comparison by the two assays, greyed out squares were removed due to low data quality.

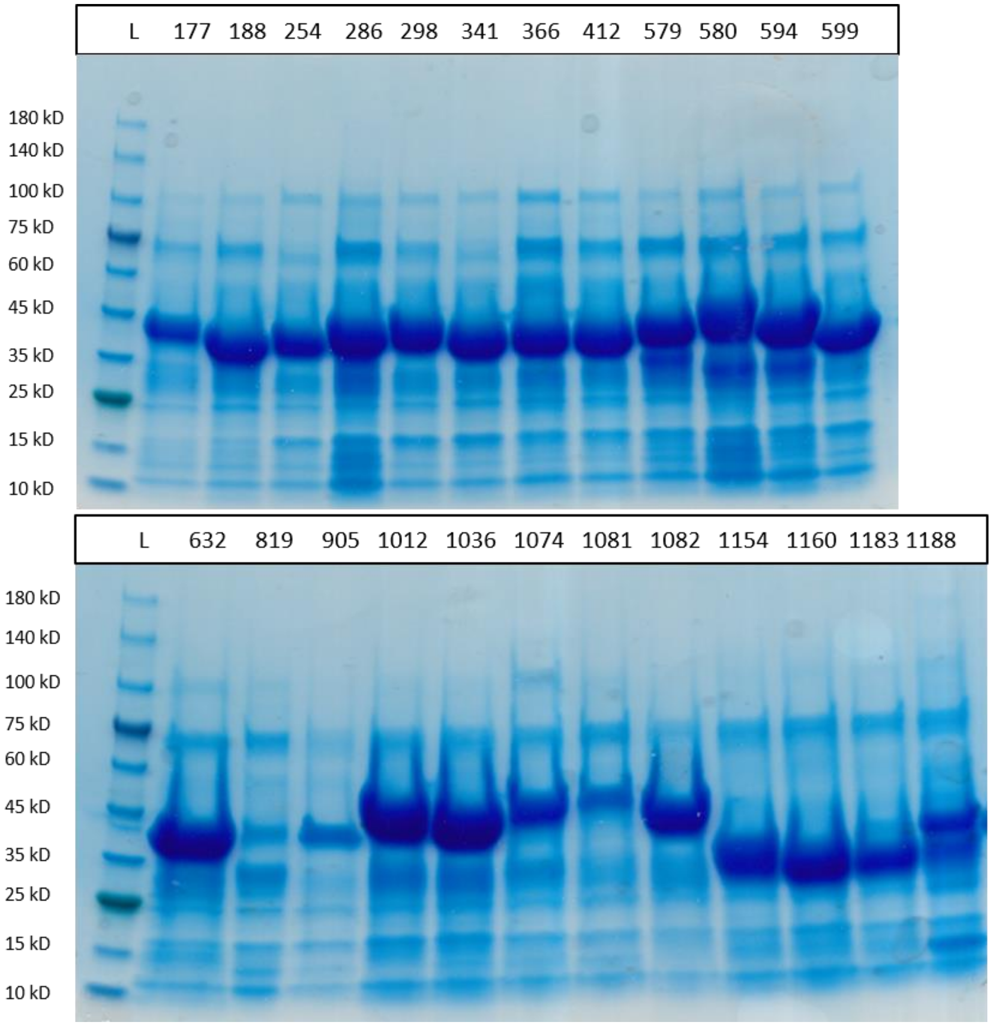


**Fig. S18.** SDS-PAGE analysis of the 24 proteins expressed for the pull-down assays. L is the protein ladder, and the numbers correspond to the Protein IDs found in Dataset S1.

**Table S1.** Background binding of EGFP to each of the substrates screened in this study.

| SUBSTRATE | AVERAGE EGFP BINDING |
|---|---|
| AVICEL | 0.13 |
| CHITIN | 0.11 |
| STARCH | 0.09 |
| PET | 0.13 |
| LDPE | 0.13 |
| PS | 0.14 |

**Table S2.** Accession numbers and origin of the catalytic domain and CBMs used to make the fusion enzymes in this study.

| PROTEIN | ORGANISM | ACCESSION NUMBER |
|---|---|---|
| $LCC^{ICCG}$ | Unknown | USU85609 |
| *RS*CBM2 | *Rivularia spp.* | AFY57603 |
| *CL*CBM3 | *Cellulosilyticum lentocellum* | ADZ83685 |
| *SD*CBM10 | *Saccharomyces degradans* | ABD81896 |
| *FL*CBM64 | *Fulvivirga ligni* | UII23919.1 |
| LINKER | *Trichoderma reesei* | P62694 |

**Legends for Datasets S1 to S4**

**Dataset S1.** Full data for CBMs selected for expression and holdup assay in this study.
Tab 1: Accession numbers and sequences for the full list of 1210 proteins designed and requested for expression by JGI. Short name codes used throughout the study are found in the column 'Protein ID'.
Tab 2: All sequences successfully synthesized by JGI.
Tab 3: All CBMs successfully expressed and purified to an acceptable concentration for use in the holdup assays.
Tab 4: Results of the holdup assays. NB: CBM did not bind to that substrate with a BI higher than EGFP alone. BD: Data for the holdup assay discarded due to bad data quality.
Link to data: https://figshare.com/s/5031fac52a7d82ee9b06

**Dataset S2.** Raw data for all holdup assays. Each 96-well microtiter plate of purified EGFP-CBM proteins tested on a single substrate is in an individual file, named with the number of the purified plate followed by the substrate.
Tab 1: All raw data containing the Protein IDs tested, the mCherry and the EGFP fluorescent signals.
Tab 2: The calculated BI of each triplicate measurement, with data filtered for quality.
Link to data: https://figshare.com/s/4267d6f858372d0602a2

**Dataset S3.** PDF files of the surface maps of the electrostatic potential and Wimley-White hydrophobicity of each CBM selected for synthesis by JGI. Four reference maps showing the location of the binding residues in each CBM family are also included.
Link to data: https://figshare.com/s/d141de1311e75f11d1ab

**Dataset S4.** The raw data for the analysis of expression of the HTP expressed proteins by LabChip GXII Calliper instrument.
Link to data: https://figshare.com/s/14422b990a095eec7510

**Supplemental Materials and Methods**

**Substrates.** All the synthetic plastic substrates used in this study were obtained from Goodfellow (UK). Commercial crystalline PET powder (PN: ES30-PD-000132) was used as purchased, while PS (PN: ST31-FM-000125) and LDPE (PN: ET31-FM-000170) were produced in-house from commercial films. Films were cut into small fragments, which were then ground in a CryoMill (Retsch, Germany) over three cycles of 1 minute at a shake frequency of 30 $s^{-1}$, with 1 min of cooling between each cycle. Avicel Ph-101(Sigma Aldrich, Germany) was used as a model substrate for cellulose. Starch granules from barley were obtained from Merck, and chitin from shrimp shells was obtained from Sigma Aldrich.

**Selection of Targets for HTP Analysis.** Full sequence sets for the CAZy families (CBM2, CBM3, CBM10, and CBM64) were obtained from the CAZy database (12). Sequences from CBM families 2, 3, and 64 were then filtered to remove 70 % redundancy using the CD-HIT software (6). For CBM10, representative sequences were selected from subgroups in a phylogenetic tree. Instances of CBM10 with tandem repeats were synthesized both as individual modules and as multi-modular repeats, preserving the native domain order (3). Full protein sequences of each of the selected CBMs without the EGFP tag can be found in *SI Appendix*, Table S1.

**Design of Gene Constructs for HTP Protein Expression.** All gene constructs were synthesized as EGFP-CBM fusions with a 6xHis purification tag on the N-terminus, and a TEV cleavage site in between EGFP (Accession number: AAB02572) and the CBM. All genes were synthesized by the Joint Genomics Institute (JGI), as part of the synthesis proposal CSP-509235 (DOI: 10.46936/10.25585/60008756). Synthesized genes were cloned into a pET28 backbone and delivered as agar stabs in *E. coli* BL21(DE3) cells.

**Analysis of Expression.** The concentration of purified proteins in 50 µL fractions was determined using a PHERAstar FSX Microplate Reader (BMG Labtech), using an excitation of 575 nm and an emission of 620 nm, calculating against a purified EGFP standard, (ThermoFisher, USA). A subset of the purified proteins was assessed by quantitative capillary gel electrophoresis on a LabChip GXII (Revvity, USA) and by SDS-PAGE. Purified proteins were then diluted to 0.5 µM in HTP binding buffer with Evo200 liquid handling robot (Tecan, Switzerland), using the 96-tip pipetting head and an eight-needle pipetting arm. Any proteins that yielded a lower concentration than 0.5 µM during purification were not diluted, with any lower than 0.2 µM not taken forward to the HU assays.

**Low Throughput Expression and Purification of Proteins.** *E. coli* BL21(DE3) cells harboring a given plasmid were used to inoculate 10 mL pre-cultures of LB medium containing 50 µg/mL kanamycin or 100 µg/mL ampicillin, which were then grown overnight at 37 °C, 160 rpm. The overnight culture was then used to inoculate 750 mL of LB medium in 3 L shake flask to an OD of 0.1 and incubated under the same shaking and temperature conditions. When the culture reached an $OD_{600}$ of 0.4–0.6, the temperature was reduced to 16 °C, and expression was induced with Isopropyl β-D-1-thiogalactopyranoside (IPTG) to a final concentration of 0.1 mM and allowed to grow for 18 hours. Cells were harvested by centrifugation at 4000 g for 30 min at 4 °C, and the cell pellet was stored at -20 °C until purification.
The stored pellet was resuspended in low-throughput (LTP) lysis buffer (10 mM HEPES, 500 mM NaCl, 10 mM imidazole, pH 7.5) and lysed by sonication, before centrifugation at 40,000 g for 30 min at 4 °C. For the purification of LCC-CBM fusion proteins, the supernatant was filtered through a 0.45 µm filter and loaded on a His-Trap FF 5 mL column (Cytiva, USA) using a peristaltic pump. The column was washed with 10 mL LTP lysis buffer, and protein was eluted with LTP lysis buffer containing 400 mM imidazole. The protein was further purified using a 16/60 75 pg Superdex column (Cytiva, USA) and a gel filtration buffer (10 mM HEPES, 150 mM NaCl, 10 % glycerol, pH 7.5). For the purification of EGFP-CBM fusions to be used in benchmarking the HTP holdup assays, the supernatant from cell lysate centrifugation was added to 5 mL of Ni-NTA beads (Cytiva, USA) and gently shaken at 4 °C for 45 minutes. The lysate was then added to a 50 mL gravity filtration column, and the beads were washed and protein eluted in 500 µL fractions of HTP binding buffer (50 mM Tris-HCl, 500 mM NaCl, pH 8.0) with increasing imidazole concentrations (10 mM, 50 mM, and 500 mM), with protein eluting in the final fraction. Imidazole was removed by dialysis in HTP binding buffer. The purified protein was stored at -20 °C. Prior to assays, the protein concentration was determined using a NanoDrop (Thermofisher, USA).

**Differential Scanning Fluorimetry Analysis.** Nano differential scanning fluorimetry (nanoDSF) was used to assess protein thermal stability in assay buffer. The analysis was done in the HTP binding buffer, with protein concentrations of approximately 0.5 µM using a Prometheus Panta (NanoTemper Technologies, Munich, Germany) at a heating rate of 1 °C per minute with a linear thermal ramp from 20 °C to 105 °C. The intrinsic protein fluorescence at 330 nm and 350 nm was measured, and the PR ThermControl Software v. 2.1.5 (NanoTemper Technologies) was used for the data analysis.
